# Optimal parameters for Rapid Invisible Frequency Tagging using MEG

**DOI:** 10.1101/2022.12.21.521401

**Authors:** Tamas Minarik, Barbara Berger, Ole Jensen

## Abstract

Frequency tagging has been demonstrated to be a useful tool for identifying representational-specific neuronal activity in the auditory and visual domains. However, the slow flicker (<30Hz) applied in conventional frequency tagging studies is highly visible and might entrain endogenous neuronal oscillations. Hence, stimulation at faster frequencies that is much less visible and does not interfere with endogenous brain oscillatory activity is a promising new tool. In this study, we set out to examine the optimal stimulation parameters of *rapid invisible frequency tagging (RFT/RIFT)* with magnetoencephalography (MEG) by quantifying the effects of stimulation frequency, size and position of the flickering patch. *Rapid frequency tagging (RFT)* using flickers above 50 Hz results in almost invisible stimulation which does not interfere with slower endogenous oscillations; however, the signal is weaker as compared to tagging at slower frequencies so the optimal parameters of stimulation delivery are crucial. The here presented results examining the frequency range between 60Hz and 96Hz suggest that RFT induces brain responses with decreasing strength up to about 84Hz. In addition, even at the smallest flicker patch (2°) focally presented RFT induces a significant oscillatory brain signal at the stimulation frequency (66Hz); however, the elicited response increases with patch size. While focal RFT presentation elicits the strongest response, off-centre presentations do generally mainly elicit a measureable response if presented below the horizontal midline. The results also revealed considerable individual differences in the neuronal responses of to RFT stimulation. Finally, we discuss the comparison of oscillatory measures (coherence and power) and sensor types (planar gradiometers and magnetometers) in order to achieve optimal outcomes. Based on our extensive findings we set forward concrete recommendations for using rapid frequency tagging in human cognitive neuroscience investigations.

## Introduction

Visual stimulation by periodic luminance or contrast change (i.e. intermittent photic stimulation) is long known to elicit an electrophysiological signal (Adrian & Matthews, 1934), the so-called steady-state visual evoked potential (SSVEP). The SSVEP frequency is aligned with the frequency of the visual stimulation and can be readily detected with intracranial as well as non-invasive electrophysiological methods such as electroencephalography (EEG) and magnetoencephalography (MEG). Rhythmic luminance changes, e.g. flickering light, are of clinical relevance across various health conditions (for review, see Vialatte et al., 2010), first and foremost in photosensitive epilepsy (Walter et al., 1946; Zifkin & Kasteleijn-Nolst Trenité, 2000), but also schizophrenia (Brenner et al., 2009), Alzheimer’s Disease (Valenti, 2013), and Parkinson’s Disease (Langheinrich et al., 2000). It has also been a valuable tool in cognitive neuroscience where specific objects on the visual display are frequency tagged, for instance when studying attention (Ding et al., 2006; Gulbinaite et al., 2019; Kim et al., 2007; Morgan et al., 1996), bistable visual perception (Chholak et al., 2020; Parkkonen et al., 2008) and brain-computer interface implementations (Middendorf et al., 2000). Originally it was thought that SSVEP in general is unaffected by attention change (Regan, 1977), but empirical findings have since demonstrated that the SSVEP amplitude is indeed robustly modulated by attention (Keitel et al., 2013; Müller et al., 2016; Müller & Hillyard, 2000; Russo et al., 2003). In particular, the magnitude of the frequency-tagged signal is thought to reflect neuronal excitability (Zhigalov et al., 2019).

From a neurophysiological perspective, periodic luminance changes might entrain cortical oscillators at the specific stimulation frequency (Herrmann, 2001; Thut et al., 2011; Zoefel et al., 2018) in particular if they target the resonant frequencies of the endogenous oscillators (Gulbinaite et al., 2019; Notbohm et al., 2016; Spaak et al., 2014) but such entrainment is not necessarily achieved in all studies (see Duecker et al. 2020). An alternative account to oscillatory entrainment by photic stimulation is that the SSVEP is merely a superposition of individual visual-evoked potentials (Capilla et al., 2011) and that the neuronal responses simply follow the stimulus with no strong entrainment of endogenous oscillators (Keitel et al., 2014, 2018). It should be noted that both entrainment and superposition of evoked responses can co-occur and the specifics are likely to depend on the experimental details.

Frequency tagging has in the past typically been applied at frequencies below 30Hz, even though it has been shown that full visual field flicker can have detectable responses up to a frequency of 100Hz (Drijvers et al., 2021; Duecker et al., 2020; Gulbinaite et al., 2019; Herrmann, 2001; Seijdel et al., 2022). Frequency stimulation below 30Hz typically produces strong responses; however, the periodic luminance changes are visible and strongly modulated by spatial attention. Furthermore, they interact with endogenous oscillations and they are likely to draw attention. Rapid frequency tagging above 50 Hz is less visible and intrusive as the drive is above the flicker-fusion frequency (Andrews et al., 1996). Furthermore, the stimulation is less likely to interfere with ongoing endogenous low-frequency oscillations in a phasic manner. These elements make rapid frequency tagging a potentially very valuable tool in cognitive neuroscience for e.g. examining attention, as well as visual processing and numerous associated dysfunctions. The main disadvantage of rapid frequency tagging is the relatively lower amplitude of the induced signal which, however, is partly mitigated by a lower noise level at higher frequencies. Therefore, it is essential to investigate which stimulation parameters result in an optimal response to achieve an ideal balance between sufficient signal-to-noise ratio and reduced visibility of the flicker.

Here we examined high frequency (60Hz and above) rhythmic luminance changes, i.e. Rapid invisible Frequency Tagging (RFT) in a three-session experiment whereby we systematically modulated key stimulation parameters and recorded the responses with magnetoencephalography (MEG, magnetometer and gradiometer sensors). In the first session, we aimed to quantify the brain response as a function of the frequency of the RFT signal. In the second session, the relationship between the brain response and the size of the flickering patch was examined. In the third session, we examined the retinotopic characteristics of the RFT by flickering different parts of the display (see Figure 1d).

**Figure 1.**
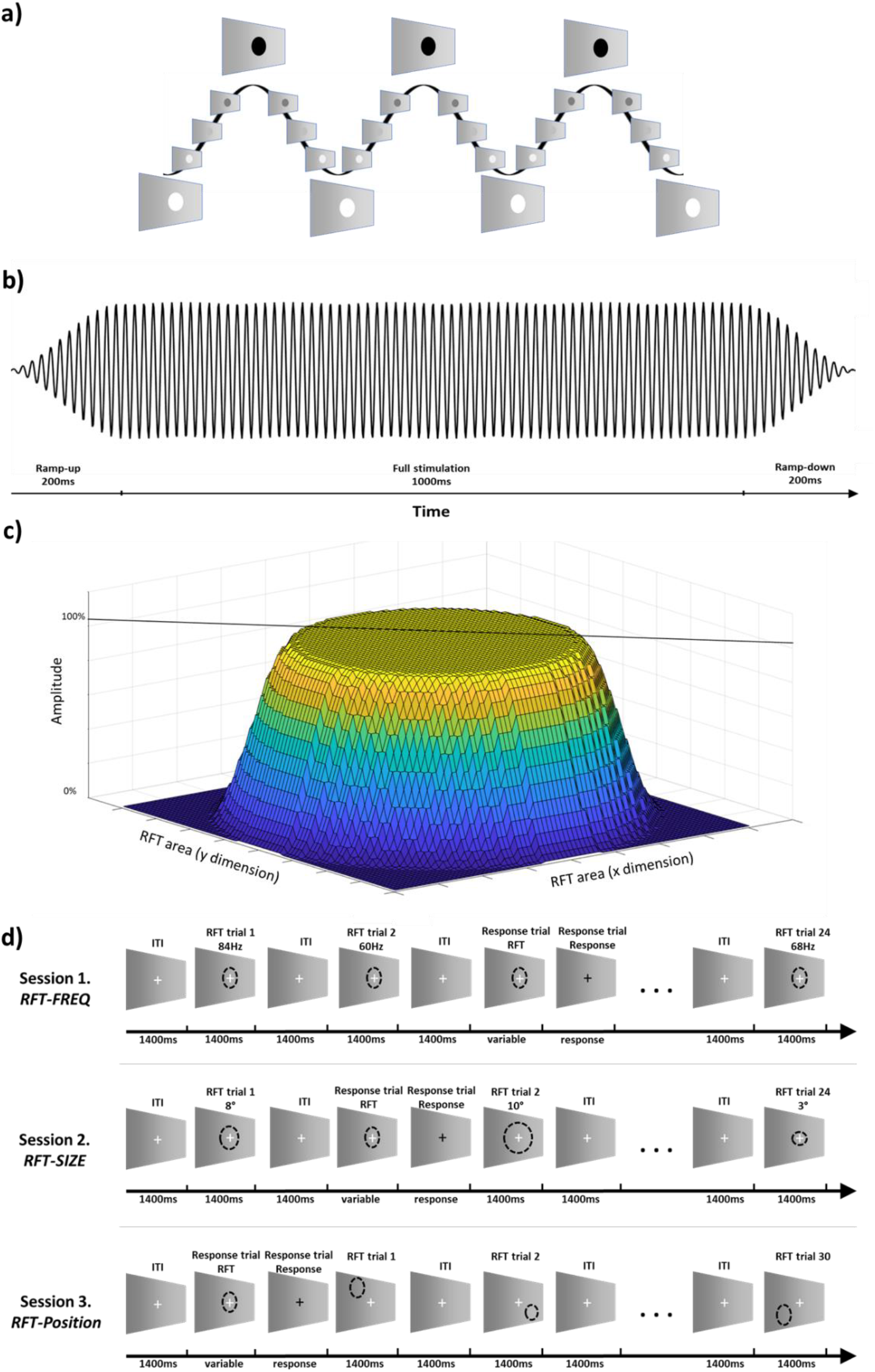
RFT stimulation and paradigm. (a) Sinusoidal modulation was applied to modulate the luminance of the stimulation patch over time. (b) The 1.4 s RFT stimulation was tapered with a 200ms ramp-up and down. (c) Spatial tapering was applied at the edges of the stimulation patch. (d) Illustration of the paradigms employed in three sessions in which RFT as a function of frequency (RFT FREQ Session 1),size of patch (RFT SIZE Session 2) and location (RFT POSITION Session 3) were investigated.

## Methods

### Participants

Twelve participants (9 female) aged between 18 and 35 years were recruited. All participants had normal or corrected-to-normal vision and were screened for participation in the MEG and MRI experiments. The participants gave written informed consent and received monetary compensation for their participation. The study received ethical approval from the Ethics Committee of the University of Birmingham and was conducted in accordance with the Declaration of Helsinki.

Data of one participant was excluded from analysis due to poor data quality and dominating heart and movement artifacts and so the final sample contained eleven participants (8 female).

### RFT stimulation

Visual stimulation was delivered using a Propixx DLP LED projector (VPixx Technologies Inc., Canada) allowing for a 1440Hz refresh rate in greyscale mode. The RFT stimulation was achieved by luminance change - effectively from black to white - following a sinusoidal pattern at a circular patch of the screen (Figure 1a). The frequency of the RFT in each condition was well above the flicker fusion threshold and therefore appeared as continuously grey, matching the background colour making the stimulation patch nearly invisible. Each train of stimulation lasted for 1.4 s. To avoid sudden onset and offset flashes at the beginning and the end of the stimulation train half Hanning tapers were applied to the input drive ensuring a 200ms ramp-up and ramp-down (Figure 1b). Thus full stimulation lasted 1 s. In addition, to attenuate a potentially visible ring at the edge of the RFT patch, the edge was spatially tapered (Figure 1c). This was done by applying a 2D taper such that full signal amplitude was delivered in a circular patch with 60% of the diameter of the total stimulation patch diameter, whereas the stimulation amplitude was tapered in the surrounding ring of 40% diameter.

### Procedure

Participants completed a visual detection paradigm combined with RFT in three separate sessions (Figure 1d) where we manipulated frequency (RFT-FREQ), size (RFT-SIZE) and position (RFT-POSITION) of the TFR patch. In each experimental session, trials requiring responses were randomly added to each block. The task was to respond to colour change (white to black) of the fixation cross. These response trials were composed of a 1.5 s inter-trial interval with no RFT stimulation, followed by a random 1 – 2s interval of RFT stimulation (see “response trial” in Fig. 1d), after which the white fixation cross turned black until response or a maximum of 500ms. Participants were asked to respond as quickly as possible by pressing the response button and received immediate feedback displaying their response time (or if they had been slower than 500ms). Response trials (25%) were randomly interspersed among regular trials (75%, see below) in each block. RFT trials were composed of a 1.6 s inter-trial interval with no stimulation and a 1.4 s RFT stimulation.

#### RFT-FREQ

The experimental session was composed of 12 blocks. In each block, 33 RFT stimulation trials and 11 response trials were completed. A stimulation patch was displayed at the centre of the screen with 6 visual degrees full stimulation size (and 10 degrees total stimulation size including the tapered ring). The frequency of the RFT stimulation in each trial was pseudo-randomized at a frequency between 60 and 100Hz in 4Hz increments. RFT stimulation was performed three times at each frequency in each block resulting in 36 trials per frequency in the full session. Note the 100Hz stimulation condition was not included in the analysis, the frequency being a harmonic frequency of the power line frequency (50Hz).

#### RFT-SIZE

The number of blocks, RFT and task trials were identical to RFT-FREQ. The size of the RFT patch ranged from 2 to 12 visual degrees sizes in 1-degree increments. Note that the specified RFT area includes the full stimulation as well as the tapered area (Figure 1c). Each patch size was used three times in each block. The presentation of each condition (patch size) was pseudo-randomized and the RFT frequency was kept 66Hz. This left a total of 36 trials per patch size.

#### RFT-POSITION

The session was composed of 16 blocks of 45 trials of which 30 were RFT trials. The size of the stimulation patch was 10 visual degrees of which the outer 4 degrees were tapered (see Figure 1c) resulting in 6 degrees of maximum stimulation. The stimulation was performed at different locations on the screen: respectively five positions above, below and in line with the fixation cross. The RFT full stimulation area of each position had no overlap with neighbouring full stimulation areas. Each position was repeated twice in each block, leaving 32 trials per location.

The screen was positioned at 100cm from the participants’ eyes. Responses were acquired using an MEG-compatible button box (Nata Technologies, Canada). The stimulus presentations were programmed in Matlab (The MathWorks Inc., Natick, Ma, US) using the Psychophysics Toolbox (Brainard, 1997).

### MEG and MRI data acquisition

MEG data were acquired with a Neuromag TRIUX (Elekta Neuromag, Helsinki, Finland) system with 306 sensors (102 magnetometers and 204 planar gradiometers). Four Head-Position Indicator (HPI) coils were placed on the participants’ heads. As part of the recording cardinal landmark points, additional scalp locations (>200 points) and HPI coils were digitized using a Fastrak (Polhemus, Colchester, USA) digitization device. Ocular artefacts were monitored and recorded with bipolar vEOG and hEOG electrodes. In addition, eye movements were also monitored with an Eyelink 1000 Plus (SR Research Ltd.) eye-tracker. Participants’ head positions were recorded once at the beginning and throughout the experiment via HPI coils. The RFT signal was recorded separately with a photodiode placed in the corner of the backlit screen.

MEG data were sampled at 1000Hz after applying a 330Hz anti-aliasing lowpass filter and a 0.1Hz highpass filter. Before each MEG recording, a 3 minute empty room MEG recording was performed. T1-weighted high-resolution 3D anatomical MRI images were acquired at the Centre for Human Brain Health (CHBH), University of Birmingham with a Siemens Prisma 3T scanner with an MP-RAGE sequence (sagittal orientation, 256×256 acquisition matrix, 208 slices, isotropic 1mm resolution). Three of the participants were previously scanned at the Birmingham University Imaging Centre (BUIC) with a Philips Achieva 3T MRI system (sagittal orientation, 256×256 acquisition matrix, 176 slices, isotropic 1mm resolution).

### MEG preprocessing and data analysis

All steps of data preprocessing and analysis were performed using the Matlab toolbox Fieldtrip (Oostenveld et al., 2010).

#### Preprocessing

The acquired MEG data were segmented in 2.8 s long trials (see Figure 1d). Each trial consisted of 1.2s baseline with fixation, 1.4s RTF stimulation and 0.2s post-stimulation fixation time window, The RFT stimulation itself consisted of 0.2s ramp-up, 1s of full stimulation and 0.2s ramp-down (see section RFT Simulation above). Task trials were not included in the analysis.

Signal space projection vectors (Ilmoniemi, 1997) were obtained from the empty room recordings for every participant individually for planar gradiometers and magnetometers via external functions (Matti Hamalainen, MGH Martinos Center, Harvard University) as implemented in the Matlab toolbox, Fieldtrip. The SSP projections were applied to the epoched data to attenuate environmental magnetic artefacts. Trials with large variance were removed from the data along with flat and noisy sensors using a semi-automatics approach. Blinks, horizontal eye-movements, cardiac artefacts and other consistent physiological artefacts were attenuated by Independent Component Analysis (Comon, 1994; Vigario et al., 2000) using the ‘runica’ algorithm (Amari et al., 1995; Bell & Sejnowski, 1995), after the data were downsampled to 300Hz. The artifactual ICA components were removed from the non-down-sampled data. Finally, the data were inspected manually and trials with remaining artefacts were removed considering both planar gradiometers and the magnetometers. The photodiode data measuring the screen flickers were segmented together with MEG trials. For sensor-level analysis, data from removed sensors were replaced by a spherical spline interpolation method (Perrin et al., 1989).

#### Power spectrum and Coherence

The analysis focused on the power spectra and time-frequency representations of the MEG signals as well as the coherence between the photodiode signal and the MEG signal.

Coherence was computed as

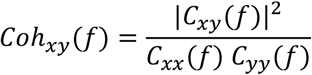

where *C*_*xy*_*(f)* is the cross-spectra at frequency *f* between signal x(t) and y(t). When *x=y, C*_*xy*_*(f)* is the power spectrum of *x(t)*. Coherence values are between 0 and 1, the latter representing the perfect alignment of the two signals.

#### Sensor-level analysis

The trials were further segmented into a baseline (−1.2 and -0.2 s, i.e. the second before the onset of the RFT ramp-up) and an RFT interval (0 to 1 s, i.e. full stimulation). The coherence and power spectra were calculated for the 1 s baseline and RFT intervals for each sensor. A 1 s Hanning taper was applied before calculating the discrete Fourier transform. Time-frequency representations over time for coherence and power were calculated between 2 and 100Hz in steps of 2Hz using a 500ms sliding time window shifted in 50ms steps. A 500 ms Hanning taper was applied to each time window prior to the spectral estimates. For the planar gradiometers, power and coherence were combined for the pairs of orthogonal gradiometers with the same locations by calculating the sum for the power and the average of the coherence.

#### Statistical analysis

The group analysis of the sensor results was performed applying non-parametric cluster-based permutation tests (Maris & Oostenveld, 2007). Pairwise comparisons involved clustering-statistics was performed using a Wilcoxon signed-rank test. The permutation p-value was calculated using 1000 random permutations. The alpha-level was set to 0.05.

To estimate the proportion of participants showing an RFT response in each of the examined conditions, each participant’s mean coherence was estimated over selected posterior sensors separately. Next, the baseline and RFT trial labels were shuffled and the mean coherence was calculated. This permutation was repeated 1000 times and a participant’s mean coherence was considered as significantly higher than chance if it was higher than 95% of the permuted mean coherence (p<.05).

#### Source analysis

To identify the source of the brain activity in response to RFT we employed Dynamic Imaging of Coherent Sources (DICS) beamforming (Gross et al., 2001). The anatomical MRI scan was co-registered with the MEG data and the anatomical volume was segmented identifying brain tissue voxels. The head model was created by a single-shell approach using spherical harmonics to fit the model to the individual MRIs (Nolte, 2003). This was then warped to an MNI template brain (Holmes et al., 1998) to align the grid points of the discretized individual brain volume to MNI space. The lead field was calculated for these gridpoints using 6mm spacing. The spatial filters were then calculated with the DICS algorithm for coherence and power of the RFT time window for all grid points and the source estimates were computed. Before plotting the source estimates were interpolated onto the template anatomical scan.

## Results

The experimental sessions were specifically designed to study the brain response to rapid frequency tagging (RFT) and to investigate how various parameters of the stimuli affect this response to optimize future studies utilising RFT in cognitive neuroscience. The following analysis is mainly based on planar gradiometer data; however, magnetometer data will subsequently be discussed and results of the two sensor types will be contrasted.

### The tagging response at 60 Hz and sensor selection

As a first step as part of the first experimental session we wanted to identify the topography and sensors of interest for the RFT stimulation. The sensor-level analysis was conducted on planar gradiometers and magnetometers separately. A cluster-based permutation test of coherence between the gradiometers and photodiode recordings revealed significantly stronger coherence (p<0.05, two-tailed) during the RFT interval with 60Hz stimulation compared to the baseline interval (Figure 2a). This difference was observed in a posterior cluster. The sources of the brain activity in response to 60 Hz RFT were localized in the striate and extrastriate visual cortex (Figure 2b). The coherence was somewhat stronger in the right hemispheric visual areas, with the maximum coherence in early visual cortical region (BA18, MNI: 8, -96, 4).

**Figure 2.**
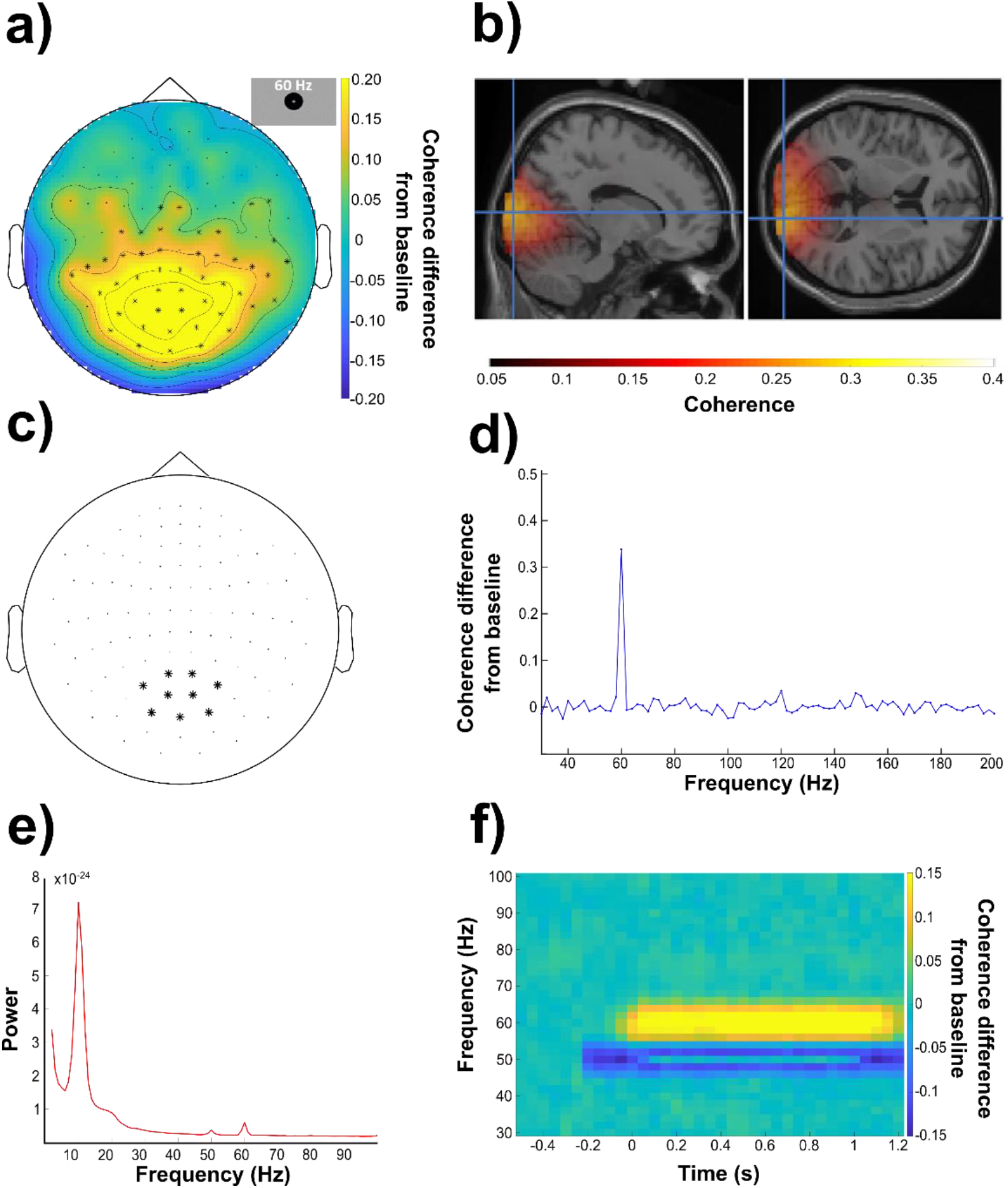
The frequency tagging response at 60Hz. (a) Topography (combined planar gradiometers) of the grand average coherence difference between stimulation and the baseline interval at 60Hz. We observed a significant difference (p<.05; cluster permutation test) over posterior regions as shown by the marked sensors. (b) Grand-average of the source localization results using DICS beamforming of the 60Hz coherence in the frequency tagging interval warped onto the MNI template brain. (c) Selected posterior sensors (combined planar gradiometers) were used in the subsequent analyses (sensors-of-interest). (d) Coherence difference (grand-average) between the interval of the 60 Hz stimulation and the baseline intervals averaged over the selected sensors of interest. (e) Power-spectrum (grand average) during the stimulation interval averaged over selected posterior sensor. (f) Coherence difference (grand average) over time during stimulation at 60Hz minus the baseline (−0.6 to -0.4s) as measured as the mean coherence over the sensors of interest.

We then focused the analysis on a selected posterior sensor set (Figure 2c). The increase in grand-average coherence (stimulation versus baseline interval) over the posterior sensors of interest showed a narrow peak at the 60 Hz stimulation frequency (Figure 2d). This peak demonstrated a robust coherence increase relative to the baseline, without obvious peaks at the higher harmonic frequencies (120 and 180 Hz). The cluster permutation test performed on the coherence at the 2^nd^ and 3^rd^ harmonics (120Hz and 180Hz, respectively) of the 60Hz stimulation did not reveal any significant clusters at either.

A cluster-based permutation test performed on the coherence calculated with a sliding interval indicated that the 60Hz stimulation effect is statistically significant (p<.05) from baseline over the 1s stimulation window (Figure S1). This is further supported by the grand-average time-frequency coherence plot relative to baseline (−0.5s to -0.2s) over selected posterior sensors (see Figure 2c for sensor selection).

### The tagging response as a function of frequency

This first experimental session was designed to assess the brain response to the flickering visual stimuli of frequencies from 60 to 96Hz (in 4Hz steps). To investigate the response as a function of frequency of stimulation, we conducted a series of cluster-based permutation tests comparing coherence during the RFT and baseline interval (combined) planar gradiometers at the stimulation frequencies from 60Hz up to 96Hz in 4 Hz steps. These tests revealed significantly increased coherence (p<0.05, two-tailed) in the RFT interval up to 84Hz (Figure S2). The results showed that the magnitude of the coherence decreased with RFT frequency. The grand average coherence relative to baseline over the selected posterior sensors showed no apparent sub-harmonics or higher harmonics peak for any of the tagging frequencies (Figure 3a). The coherence (grand-average; selected sensors) declined linearly starting from 0.50 (SEM=0.04) at 60Hz stimulation decreasing to baseline levels at 88Hz (*M*_*tag-60Hz*_=0.19, SEM_*tag-60Hz*_=0.01, *M*_*base*_=0.17, SEM_base_=0.01; Figure S3a). A linear fit demonstrated a slope of 0.046/4Hz decrease in coherence (SEM=0.0055).

**Figure 3.**
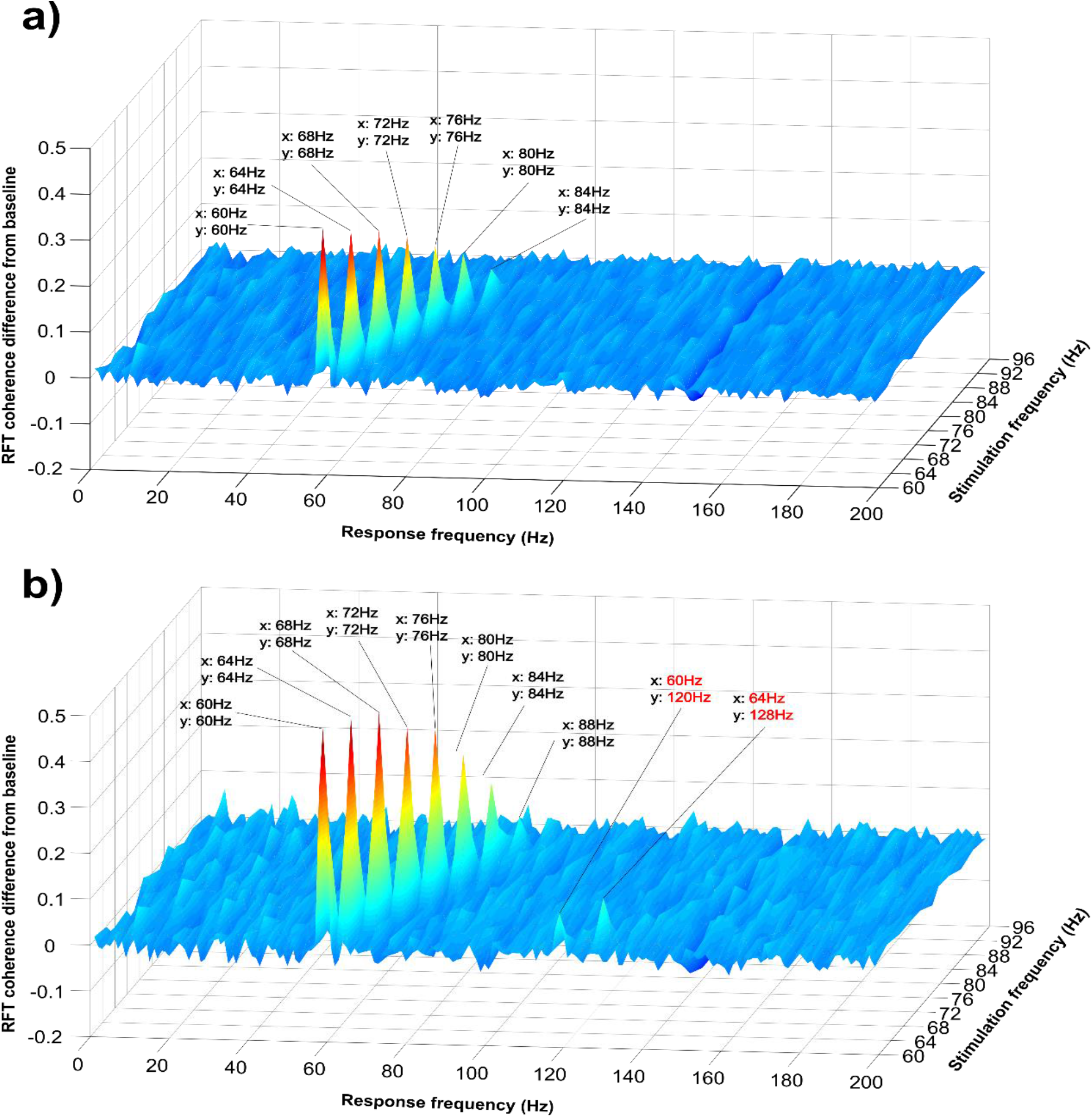
Coherence difference (grand average, z-axis) between the stimulation and baseline interval over the posterior sensors of interest as a function of stimulation-frequencies from 60Hz to 86Hz in 4Hz steps (y-axis). (a) Combined planar gradiometers. (b) Magnetometers.

Noteworthy is further that the frequency tagging-induced brain response varies across individuals (Figure 4a; S3a). When expressed as a ratio of RFT-interval coherence relative to baseline coherence at 60Hz, the highest individual score indicates 5.1 times increase of coherence, whereas the lowest individual score 1.95 times relative to the baseline level (Figure a; left). At 84Hz the highest value indicates 1.94 times increase of coherence relative to baseline, whilst the lowest score is 0.98, effectively showing no change from baseline.

**Figure 4.**
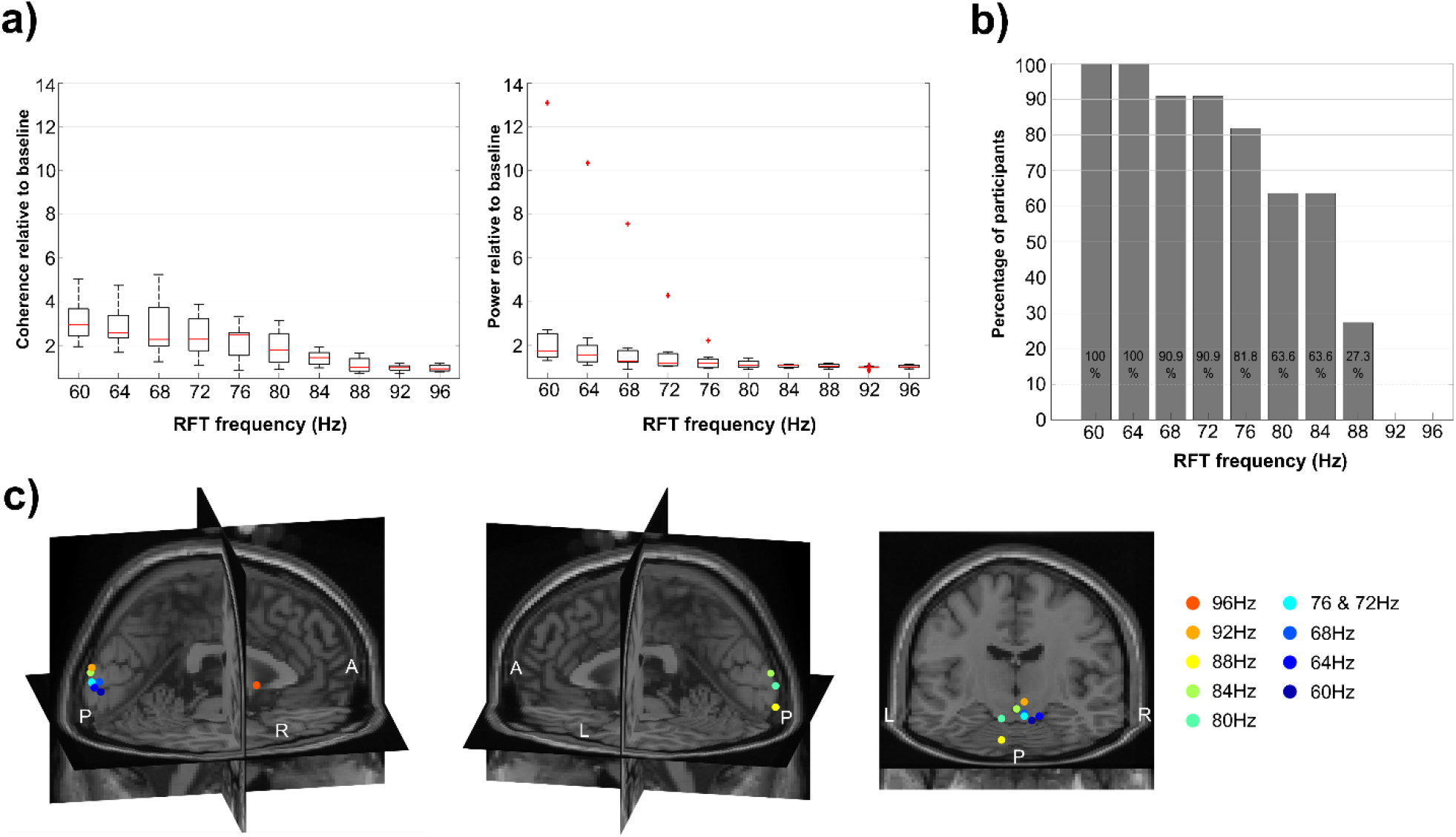
TFR examined as function of frequency. (a) Boxplot of coherence (left) and power (right) during RFT relative to baseline across conditions. Red lines indicate the median, the boxes of the 25^th^ and 75^th^ percentiles and the whiskers the minimum and maximum values without outliers. Outliers are displayed as red crosses. (b) Percentage of participants showing significant coherence difference between the stimulation interval and baseline. Percentage calculated based on permutation of mean trial coherence of the RFT and the baseline intervals over selected posterior sensor cluster of each participant. (c) Maximum of the grand-average of the source localization RFT coherence results with DICS beamforming of each condition in the stimulation interval. The results are mapped on a MNI template brain.

This variability in brain response as quantified by coherence leads to the question of how many participants actually show a significant RFT response across the conditions. This issue was assessed by permuting the trial data over selected posterior sensors for each participant (see methods). The obtained scores were used to investigate if the coherence at a stimulation frequency of a participant was significantly (*p*<.05) higher than expected by chance. The results show that at 60Hz and 64Hz all of the participants had a significant increase in coherence. However, the proportion decreases steadily with increasing frequency (Figure b). At 92Hz no single participant had a coherence higher than expected by chance.

The sources of the maxima of the RFT coherence were localized mainly to the primary and extrastriate cortex with the maximum near the occipital pole (**Error! Reference source not found**.c; see Figure S4 for the whole estimated source). The maximum distance between any of these coherence maxima source points (excluding the 96Hz condition) was 29.6 mm. Considering only the maximum points of conditions with significant cluster-based permutation test results, i.e. from 60Hz up to 84Hz, the maximum distance is 21.9 mm. As such, while there was some spread in the localization for sources, they were not systemically arranged according to frequency.

### The frequency tagging response as a function of size of the stimulation patch

The second experimental session examined how the frequency tagging response varied with respect to the size of the stimulation patch (increased in size from 2° to 12° in steps of 1° ; note that each patch was spatially tapered at the edges) at 66Hz. We performed a cluster-based permutation test for each patch size comparing the stimulation and baseline interval. This revealed a significant coherence increase for all patch sizes starting from 3° (Figure 5, Figure S5). The grand average of the posterior sensors-of-interest, reveals a coherence increases with the size of the patch (from 0.22 +/-0.02 to 0.46 +/-0.02) without showing an obvious plateau (Figure 5a left). The fitted linear slope to the grand average shows 0.023 coherence/degree increase (SEM=0.0026).

**Figure 5.**
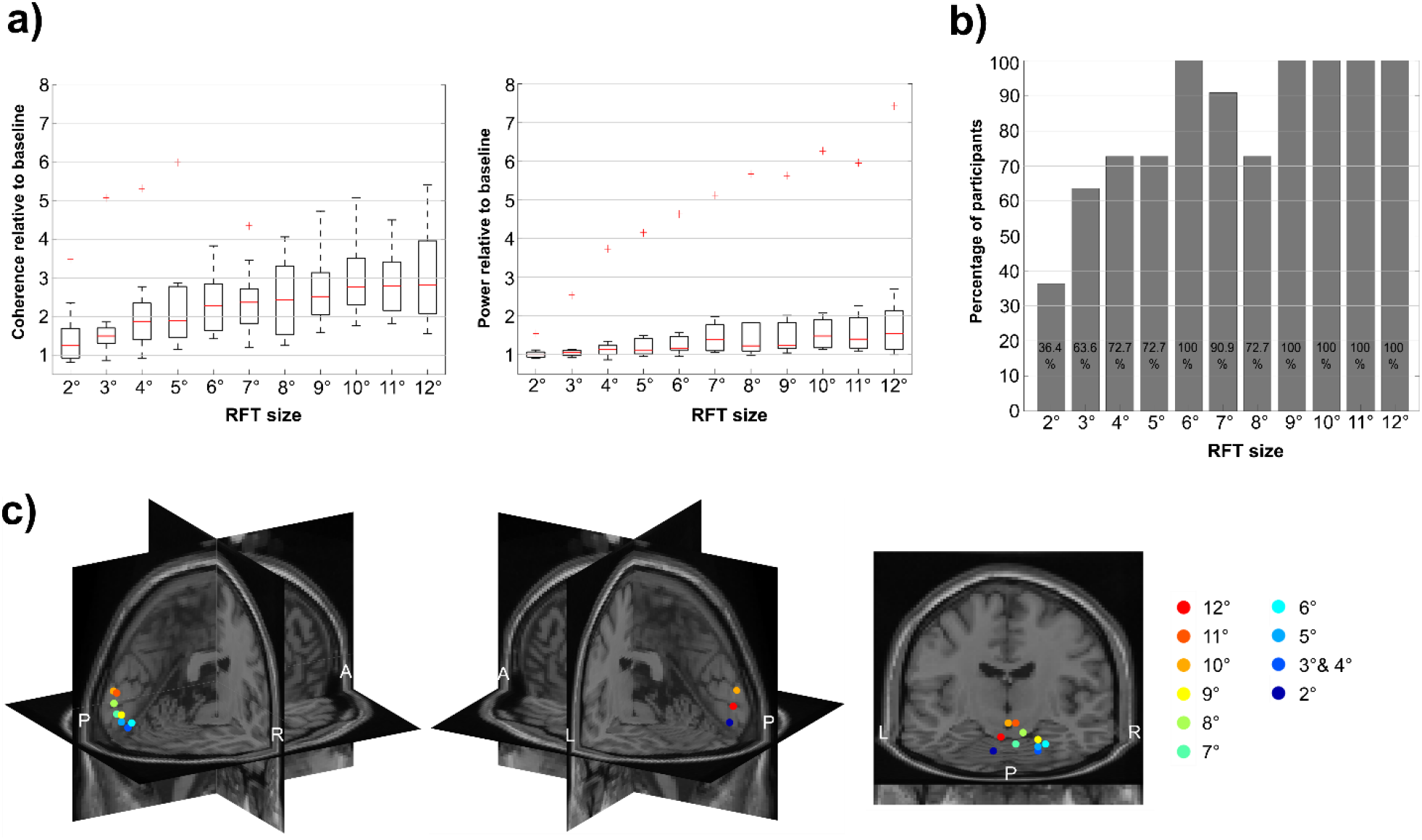
Frequency tagging response as a function of patch size. (a) Boxplot of coherence (left) and power (right) during stimulation relative to baseline. (b) Percentage of participants showing significant coherence difference between the frequency tagging and baseline interval. (c) Maxima of the localized sources using DICS beamforming for the individual patch sizes. The results were warped on the MNI template brain.

The measured brain response to frequency stimulation with patch sizes varied across individuals (Figure 5b, Figure S3d). Whilst the group statistics revealed no significant increase from baseline in the stimulation interval at 2° patch size, it is worth emphasizing that 36.4% of the participants did show a significant response (*p*<.05) in the sensors-of-interest. Group statistics were significant from 3° and proportion of participants showing a significant response display a steep rise from 5° to 6°. From 9° size upwards, all participants showed a significant increase in coherence from baseline. The participant with the highest coherence increase from baseline over selected posterior sensors had a 3.5 fold coherence increase at patch size 2°, whereas the individual with the lowest response had an 0.80fold increase. At path-size 12°, the individual with the strongest response had a 5.4-fold increase, compared to a 1.6-fold increase for the participant with the weakest response.

Source localization of the coherence with the photodiode at 66Hz indicated largely occipital and posterior cortical sources with the maximum of each RFT size condition near the occipital pole (Figure 5c, see Figure S6 for the whole estimated source). The maxima of the coherence across RFT sizes between any two conditions were within 36.2 mm.

### The frequency tagging response as a function of position

The third session was implemented to provide empirical data regarding the RFT response delivered at different parts of the visual field, i.e. patches centrally and peripherally presented on a screen with increasing eccentricity. We used a patch with the same size (10° total size including the 40% tapered area, see Figure 1c) and frequency (66Hz) across all trials but at different positions (Figure 6a & b). The cluster-based permutation test at each position indicates that in 10 out of the 15 examined positions there was a significant increase in coherence between the stimulation and baseline interval (Figure c). The grand average coherence over selected posterior sensors suggests that the RFT response is weaker for the upper visual field and for the more lateralized positions (**Error! Reference source not found**.b). To statistically assess how position affects the brain response, we conducted a repeated-measure ANOVA on the mean coherence over selected posterior sensors with the factors Horizontal (5 levels) and Vertical (3 levels) positions. The results revealed a significant main effect of Vertical position, *F*(2,20) = 42.91, *p* < .001. Pairwise comparisons indicated that the coherence was significantly larger when the RFT was delivered in the Middle than in the Upper row, (*p* = .001), and in the Lower than in the Middle row (*p* = .005), as well as in the Lower compared to the Upper row (*p* < 0.001). These results suggest a vertical direction bias, in particular, coherence increases from Upper over Middle to Lower RFT row positions. In addition, the analysis revealed a significant main effect for the factor Horizontal position, *F*(4,40) = 34.93, *p* < .001. Pairwise comparisons suggest that lateral comparisons showed no significant difference; where the Extreme Left positions versus the Extreme Right positions (*p* = 1.00) and the Left positions versus Right positions (*p* = 1.00) did not produce a RFT signal. All other position comparisons revealed a significant effect (*p* < .05); i.e the farther the position is away from the center the lower the coherence (*M*_*Extreme* *Left*_ = 0.21, *SEM* = 0.01; *M*_*Left*_ = 0.32, *SEM* = 0.02; *M*_*Central*_ = 0.42, *SEM* = 0.03; *M*_*Right*_ = 0.31, *SEM* = 0.02; *M*_*Extreme Right*_ = 0.21, *SEM* = 0.01). These results demonstrated that the coherence decreases as lateral distance from the focus of vision/attention increases, irrespective of the hemifield in which the RFT was presented. Finally, the analysis also revealed a significant interaction between the Horizontal and Vertical position factors *F*(8,80) = 5.87, *p* < .001. This was driven by the increasing coherence from the Upper Row, over the Central Row to the Lower Row, but not to the same extent for the two hemifields. The least amount of increase was observed in the Extreme Left and Right horizontal positions and most in the Central positions (Figure S7).

**Figure 6.**
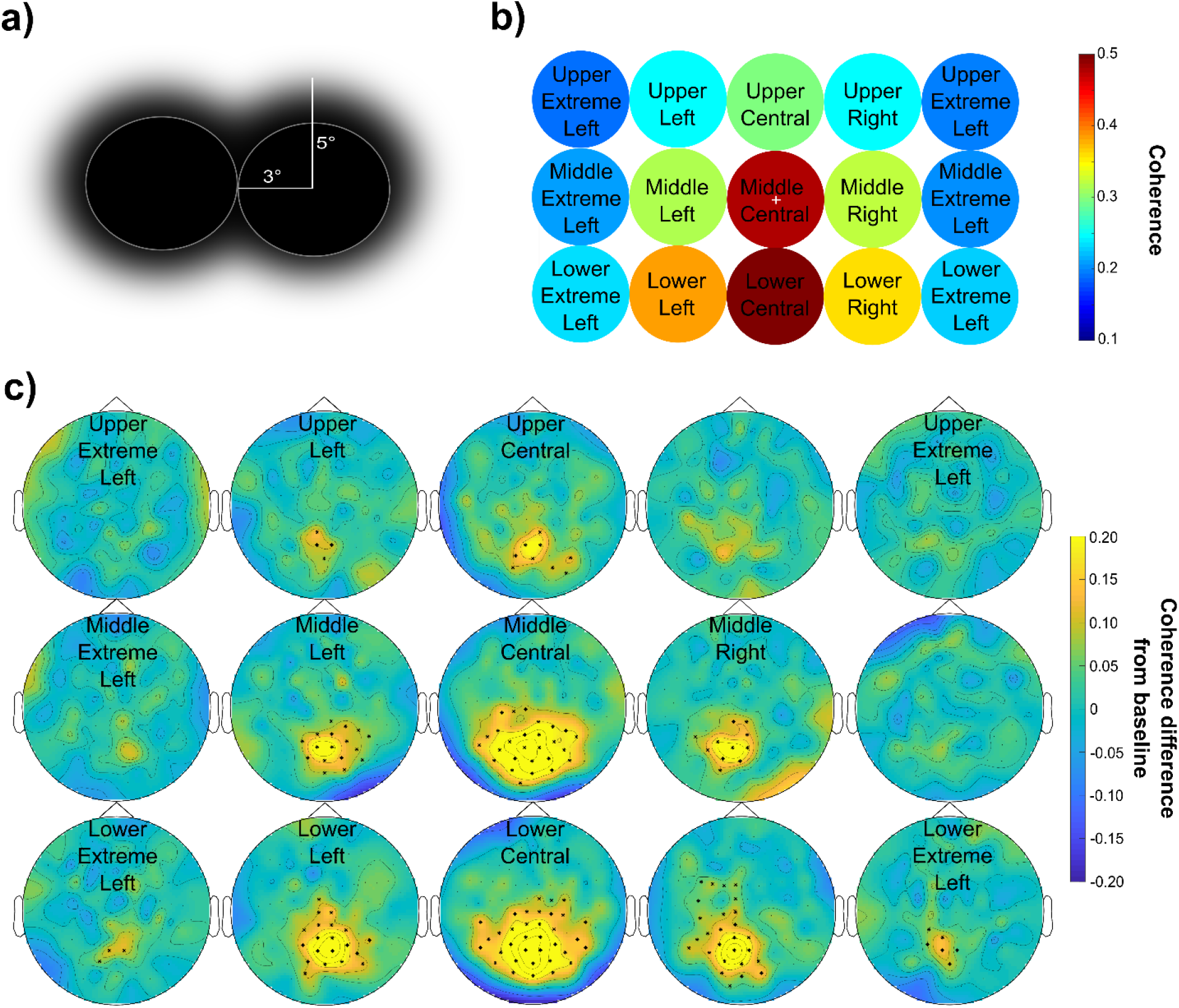
(a) Schematic depiction of the RFT area and relative position. The RFT area with full amplitude (white circle) is surrounded by a tapered area (faded area). Two neighbouring positions are aligned vertically and/or horizontally with the centres being 6° apart with no overlap between the full stimulation areas. (b) Heat map of grand average coherence over selected posterior sensor cluster for each RTF position. The position of each condition (see labels) in the plot corresponds to the position of the tagged area on the screen during the experiment. The white cross indicates the fixation cross. (c)Topography of the grand average coherence difference between RFT and baseline interval at 66Hz in response to 66Hz stimulation at each position in the RFT-POSITION session. The highlighted sensors show a significant difference between the RFT and baseline interval after performing a cluster permutation test.

In sum, these findings suggest that coherence decreases with eccentricity and with the height in visual field of the RFT stimulation.

The sources of the coherence with the photodiode at 66Hz again largely fell into the primary and extrastriatal visual cortices with a clear retinotopic arrangement (see Figure 7c, see Figure S8 for the whole estimated source) extending to other regions in conditions mainly when the coherence is larger. The largest distance of any two condition coherence maxima at the source level was at 35.7 mm.

**Figure 7.**
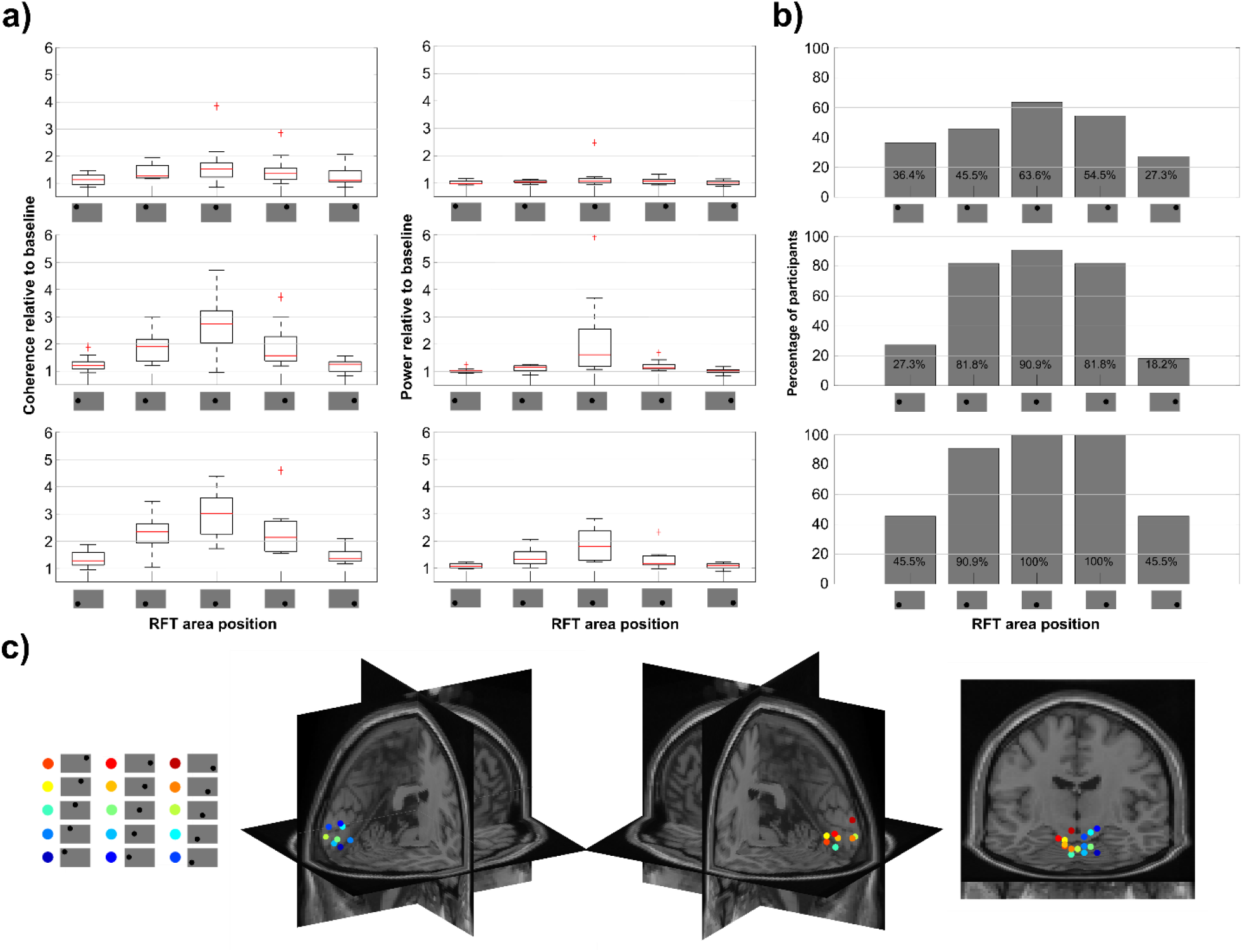
The RFT-POS condition. (a) Boxplot of coherence (left) and power (right) during RFT relative to baseline across conditions. (b) Percentage of participants showing significant coherence difference between RFT interval and baseline. (c) Maximum of the grand average of the source localization RFT coherence results with DICS beamforming of each condition in the stimulation interval. The results are warped on the MNI template brain.

### Individual variance

Participants’ measured response to RFT stimulation at various RFT locations varied considerably. At the Mid-Central location, the participant with the highest coherence had a 4.7 fold increase from baseline whereas the participant with the lowest signal showed no increase, with 0.96 coherence relative to baseline (Figure 7a). Of course, at positions with weak or no RFT effect in general, the highest and lowest individual ratios are also considerably lower. For instance, at the Upper Extreme Left position, the highest individual ratio indicates a 1.5 fold coherence increase relative to baseline whilst the participant with the lowest ratio a 0.9 coherence relative to baseline.

To establish what ratio of participants showed a significant RFT response, coherence over selected posterior sensors was calculated against a set of permuted (baseline and RFT intervals) coherence values (Figure 7b). This showed that Lower Central and Lower Right locations were the only areas where 100% of the participants’ RFT responses were significantly higher than baseline. Furthermore, Lower Left, Middle Central, Middle Right and Middle Left positions had significant coherence in more than 80% of the. participants. The Upper Central position revealed only 64% of the participants showed a significant RFT response. The percentage drops dramatically with increasing eccentricity in every row.

### Lateralization

For the analysis of hemispheric lateralized RFT presentation, the lateralization index (LI) was calculated as follows:

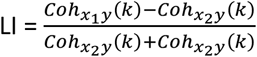

where x_1_ is a left and x_2_ a right hemifield RFT area. The frequency (k) considered was 66Hz.The previously described, statistical results indicated that the RFT effect is dependent on the position of the RFT area on the screen. However, the analysis so far did not address the question of whether there is left-right hemispheric lateralization when right and left hemifield RFT locations are compared. To address this question, a series of cluster-based permutation tests were performed with each lateralized position and its opposite hemifield counterpart contrasted. Thus, when at least one of the positions showed a significant increase in the RFT frequency coherence compared to the baseline, it and its counterpart on the opposite hemifield were fed into the cluster-based permutation test. The statistical results only showed a significant difference with a higher coherence over a small posterior right hemispheric sensor cluster for the *Lower Left* vs. *Lower Right* comparison (Figureb, left) but no significant differences (*p>*.*05*) in any of the remaining comparisons (*Upper Left* vs. *Upper Right, Middle Left* vs. *Middle Right, Lower Extreme Left* vs. *Lower Extreme Right*).

### POWER

The analysis of the RFT response so far has been focusing on the coherence between the signal recorded with the MEG and the photodiode, but we also set out to examine if RFT produces a reliably detectable brain response in more conventional and widely used time-frequency power analyses. Note that, unlike coherence, power can serve as input for trial-based analysis.

#### RFT-FREQ

To investigate if an RFT brain response can be detected by analysing oscillatory power a series of cluster-based permutation tests was conducted from 60Hz to 96Hz in 4Hz steps, identical to the analysis of coherence. The results revealed significant differences between the RFT interval and the baseline interval over a cluster of posterior electrodes from 60Hz up to a maximum frequency of 76Hz (Figure S9); which is considerably lower than the maximum frequency detectable with coherence (84Hz, see Figure S2). The grand average power plot of selected posterior sensors of the RFT interval at 60Hz indicates that the power peak is visible, but modest especially in comparison to the alpha peak around 10Hz (Figure 2e).

To further compare results obtained with power and coherence as measures, we examined the median power and coherence of the RFT time window relative to baseline over selected posterior sensors. The median coherence measured over posterior channels at 60Hz is 2.95 times (SEM = 0.27) the baseline, whereas median power is only 1.74 times (SEM = 1.03) the baseline (Figure 4a). By 88Hz, this RFT-baseline ratio of both median coherence and power indicates effectively no increase during the RFT interval relative to baseline (Mdn_coh-ratio-88Hz_ = 0.99, SEM_coh-ratio-88Hz_ = 0.10, Mdn_pow-ratio-88Hz_ = 1.04, SEM_pow-ratio-88Hz_ = 0.03). The fitted linear slope to the median coherence and power across stimulation frequencies suggests that the RFT-baseline coherence ratio decreases by 0.28/4Hz (*SEM* = 0.04) between 60Hz and 88Hz, whereas power ratio decreases by 0.10/4Hz (*SEM* = 0.16) over the same frequency interval.

A further aspect to consider is that the RFT response measured by power ranges more substantially across participants than coherence (Figure 5a and Figure S3a-c). This is largely due to one participant, whose data at 60Hz showed at least a one magnitude larger RFT response measured with power than any other participant (13.18 times of the mean baseline level).

#### RFT-SIZE

Whilst cluster permutation tests on coherence showed significant differences between RFT and baseline intervals from the 3° size upwards, the same statistical tests reveled significant differences only from 6° and above when the brain response is measured with power (Figure S10).

The relative median power change from baseline over selected posterior sensors indicates that the increase in power over the RFT interval is considerably lower across all sizes than the relative coherence change (Figure 7a). At 2° the median coherence change is 1.26 (*SEM* = 0.24), whereas the median power change is merely 1.00 (*SEM* = 0.05), meaning no change from baseline. At 12° size the RFT interval coherence is 2.81 times the baseline (*SEM* = 0.38), whereas with power it is only 1.54 (*SEM* = 0.55). The linear slope fitted to the median coherence and power relative to baseline ratio indicated that the relative change to baseline increases much faster in the case of coherence than power (*Mdn*_slope-coh_ = 0.144, *SEM*_slope-coh_ = 0.022, *Mdn*_slope-pow_ = 0.053, *SEM*_slope-pow_ = 0.042). Finally, when power is used as measure instead of coherence, one participant showed again a markedly higher RFT response (Figure S3d-f) than any other participant (*Pow-ratio*_*2°*_=1.54, *Pow-ratio*_*12°*_=7.42).

#### RFT-POSITION

When assessing RFT response measured with power as opposed to coherence, the performed series cluster-based permutation test showed only 6 out of the 15 positions with a significant difference between baseline and the RFT time-interval (Figure S11; in contrast with 10 out of 15 when measured with coherence). Moreover, comparing power versus coherence change relative to baseline at 66Hz over posterior sensors indicates that the median RFT-baseline ratio is higher when measured with coherence than power at every RFT position (Figure 7a).

### MAGNETOMETERS

The analysis so far has focused on the results of the planar gradiometers, here we will briefly discuss the findings based on magnetometers. As findings of the two types of sensors are largely similar, this section will mainly discuss their differences.

#### RFT-FREQ

The series of sensor-level cluster based permutation tests at the stimulation frequencies revealed a significant difference between baseline and RFT interval from 60Hz up to 84Hz, similarly to the planar gradiometers (Figure S12). However, the magnetometers revealed results on sensor-level with more sensors showing a significant difference than planar gradiometers. When examining the grand average coherence over a selected posterior sensor cluster (same location as the selected planar gradiometers), the magnetometers showed higher mean coherence than planar gradiometers across the entire frequency range up to 92Hz (*Figure 9*a); indicating better SNR for magnetometers in general.

Furthermore, in a series of cluster-based permutation tests we examined if there was a significant increase during the RFT interval relative to baseline in any of the second harmonic frequencies. Unlike the planar gradiometers, at 60Hz and 64Hz stimulation frequencies there was a significant increase at 120Hz and 128Hz, respectively (Figure S13) over mainly a small cluster of posterior sensors. These harmonic peaks can also be observed in the grand average over selected posterior sensors (Figure 3b).

The sources of the coherence at the stimulation frequencies are primarily occipital (Figure S14) with the exception of the 96Hz condition, similarly to the results of the planar gradiometers. The maximum of the source level coherence of each condition was near the occipital pole – again except for the 96Hz stimulation condition - with the largest distance between any two conditions’ maxima being 27.2 mm (Figure S15a). The difference between the source activity maxima localized with planar gradiometer and magnetometer data did not exceed 16.49 mm (*M* = 11.97, *SEM* = 1.06) at any of the conditions, when the 96Hz condition is not taken into account (Figure S16a). When focusing on the conditions which showed significant coherence increase from baseline with cluster permutation test (60Hz to 84Hz, in 4Hz steps), the largest difference remains the same (*M* = 12.32, *SEM* = 1.12).

#### RFT-SIZE

The performed series of cluster-based permutation statistical tests indicated that, in contrast to the planar gradiometers (3°-12°), significant differences emerged between the RFT and baseline interval at all of the examined RFT area sizes, starting already from the smallest size of 2° to 12° (Figure S17). However, at 2° the difference is over a right-central sensor-cluster, away from typical posterior sensor positions, warranting some caution in terms of interpretation. From 3° to 12°, the sensor clusters are increased and in general larger than the equivalent comparisons with planar gradiometers. Furthermore, the grand average coherence over selected posterior sensors was generally higher using data from magnetometers than planar gradiometers at every condition from 2° to 12° sizes (Figure 9b).

The source localization of the coherence showed occipital sources at each size condition (Figure S18), with source maxima near the occipital pole falling almost exclusively in the right hemisphere (Figure S15b). The largest distance between any two of these condition maxima was 25.61 mm. When comparing the coordinates of the source maxima of each condition (2° to 12°) obtained with magnetometers as compared to planar gradiometers the largest distance between the maxima obtained with the two different types of sensors was 27.20 mm across the RFT sizes (*M* = 11.33, *SEM* = 1.73; see Figure S16b)

#### RFT-POSITION

To test if there is a significant difference between the RFT and the baseline interval at sensor-level cluster-based permutation tests were performed at each RFT position. There were only two positions, Upper Right and Central Extreme Right, where the two intervals did not differ significantly (Figure S19). Moreover, when looking at the grand average coherence over selected posterior sensors it is apparent that across all positions the mean coherence is larger obtained with magnetometers than planar gradiometers (Figure 9c).

The sources of the coherence at 66Hz of each RFT position were primarily in the occipital cortex (Figure S20). The source maximum of each condition was again around the occipital pole, with the only exception of the Upper Extreme Left condition (Figure S15c) being localized up to 78.82 mm from other condition maxima. Excluding the Upper Extreme Left condition, the largest distance between any two condition source maxima was 43.95 mm. Finally, comparing the source maxima coordinates of the coherence between magnetometers and planar gradiometers showed that the largest difference was 47.03 mm (*M* = 12.33, *SEM* = 2.81), which again was due to the difference in the Upper Extreme Left condition. Without this condition, the largest distance between the source coordinates of the two types of sensors is 20.88 mm (*M* = 9.85, *SEM* = 1.43; see Figure S16c).

##### Lateralization

A series of cluster-based permutation tests were performed comparing left and right hemispheric RFT presentation including conditions whereby at least one of the RFT positions led to a significant RFT increase from baseline (*Upper Left* vs. *Upper Right, Middle Left* vs. *Middle Right, Lower Extreme Left* vs. *Lower Extreme Right, Lower Left* vs. *Lower Right*). In comparison to the planar gradiometer results, the magnetometer results strongly indicated lateralization effects. In addition to the increased coherence in a right hemispheric sensor cluster (as also obtained with the gradiometers), significantly lower coherence (*p*<.05) was found at the stimulation frequency (66Hz) over an additional left hemispheric sensor cluster when the RFT stimulation was presented at the *Middle Left* as compared to the *Middle Right* position (left minus right comparison as outlined in the section *Lateralization* for the gradiometer sensors previously). In addition, when comparing *Lower Left* and *Lower Right* RFT presentation, the cluster-based permutation test revealed a left hemispheric posterior sensor cluster with significantly lower coherence and a significantly higher coherence over a right hemispheric sensor cluster. This latter finding shows a clear lateralization effect on coherence in *Lower Left* and *Lower Right* conditions (see Figure 8, right)

**Figure 8.**
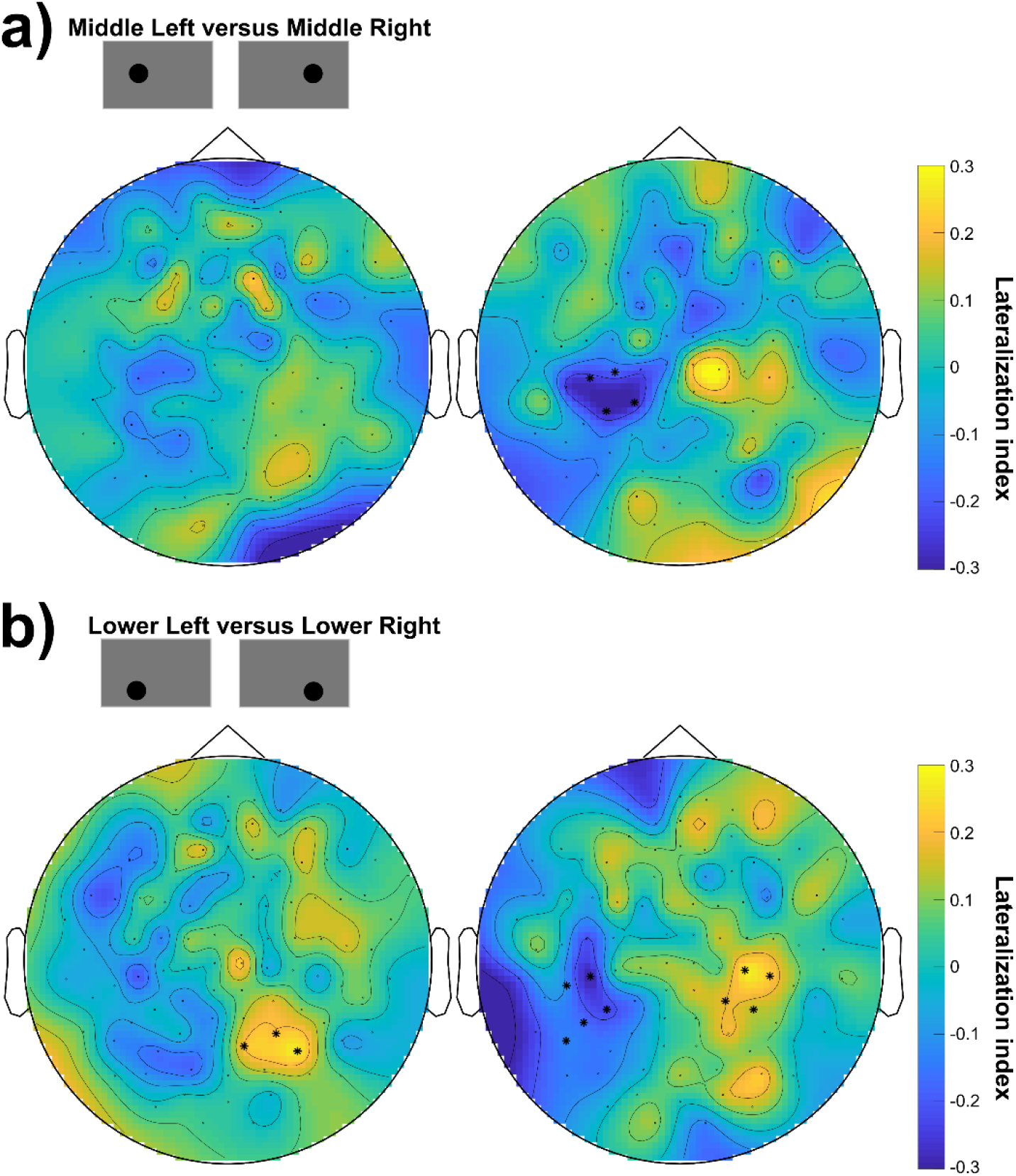
Lateralization of cortical response to lateralized RFT stimulation. (a) Condition comparison of RFT delivered at the Lower Left and Lower Right positions. Cluster-based permutation test results with highlighted sensors indicate a significant difference between the two conditions (p<.05). **Left: Planar gradiometers. Right: magnetometers**. (b) Same as (a) with the comparison Lower Left and Lower Right. Left: Planar gradiometers. Right: magnetometers.

**Figure 1.**
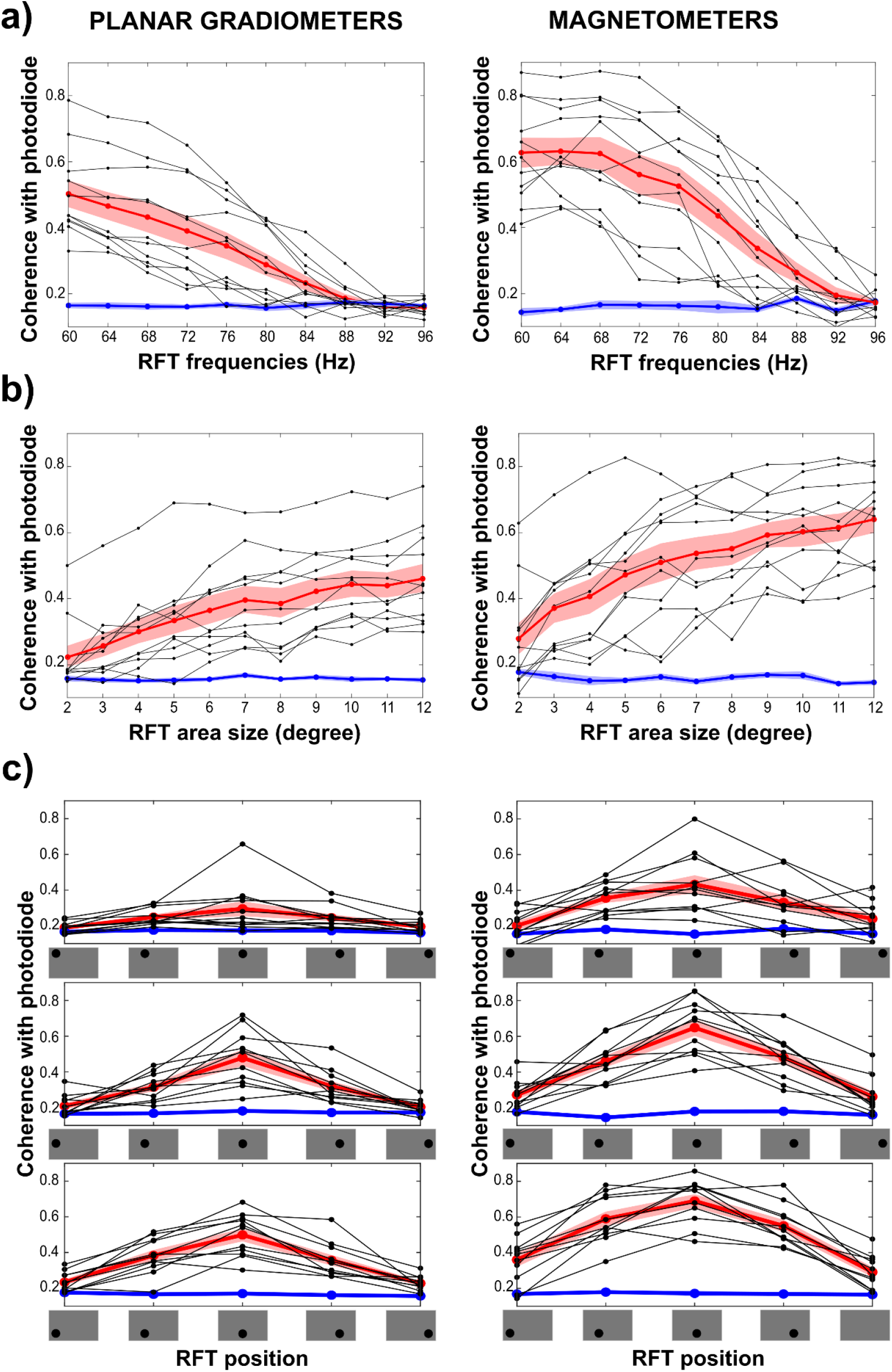
(a-c) Coherence with photodiode at the stimulation frequency in the RFT interval comparing selected posterior planar gradiometers (left) and magnetometers (right) in the (a) RFT-FREQ, (b) the RFT-SIZE and (c) the RFT-POSITION sessions. Grand average RFT interval coherence (red line), baseline interval coherence (blue line) and individual mean RFT interval coherence (black lines) are shown.

## Discussion

The findings of the study confirm that rapid frequency tagging (RFT) in the form of fast luminance changes produces a reliably and highly frequency-specific cortical response. Importantly we can put forth a number of recommendations regarding the optimal parameters of stimulation. Taking the vast experimental results into consideration we would recommend using a flicker frequency below 72Hz. The stimuli should be shown within a 9° angle and in the lower visual field. Typically the response increases closer to the midline and with the size of the stimuli. These conclusions are based on the presentation of a single stimulus delivered in the absence of other stimuli but they should generally apply to experimental paradigms utilising RFT.

### Frequency dependence

It is well established that the amplitude of the SSVEP decreases with higher frequencies. Whilst it has been reported that SSVEP can be detected up to 100Hz with full visual field ON-OFF flicker (Herrmann, 2001) our study is using a circular RFT patch with a sinusoidal signal which consistently showed a significant flicker response up to 84Hz. The grand-average coherence quantifying the flicker response showed a steady decline as the stimulation frequency increased from 60Hz to 88Hz RFT. The decrease of coherence at higher frequencies is in line with the well-documented amplitude decrease of SSVEP at higher frequencies across various stimuli (Pastor et al., 2003; Duecker et al., 2020). No participant showed a significant coherence increase with respect to baseline above 88Hz stimulation frequency. Another consideration that needs to be taken into account is that it is only the 60Hz and 64Hz stimulation frequencies where every participant showed a significant coherence increase from baseline at the stimulation frequency, when measured with planar gardiometers. Up to 72Hz about 90% shows a robust response, but above 72Hz it drops relatively quickly.

### Size

Besides the frequency of the RFT stimulation, the size of the stimulation area is also a key consideration. Even though a patch of 3° showed a significant coherence increase in the magnetometers, only 64% of the participants showed a significant RFT increase. For patches above 9° in size all participants displayed a significant RFT increase irrespective of sensor type. It also needs to be mentioned that the grand-average coherence showed a steady increase over the size range from 3° to 12° degree sizes with no sign of plateauing. In short, the larger the flickering path, the larger the response.

### Location and retinotopy

As expected the RFT response was very consistently retinotopically organized. Stimulation in the left hemifield displays a stronger response over the right hemisphere and vice versa. Patches presented above the visual midline have their maxima in the visual cortex lower than the patches presented below the visual midline. Furthermore, stimulation in the lower hemifield presentation produced a stronger response than in the upper hemifield. Secondly, the closer to the center of fixation the RFT area is presented on the horizontal plane, the stronger the RFT response. Importantly, it was only at the *Lower Central* and *Lower Right* positions where all of the participants showed a significant RFT response. The decreased coherence in response to RFT away from the focus is likely due to attentional gain, but the vertical bias may be because of the differential retinotopic representation of upper and lower visual field stimuli. Since the upper visual field is represented to a large extent deeper down the calcarine sulcus, whereas the lower visual field dorsal to it (Wandell et al., 2005), MEG sensors are likely to be more sensitive to activity in the latter area due to sensore proximity (see also Portin et al., 1999).

### Source modelling

The source localization results across the various conditions indicating the RFT response was constrained to occipital cortex. Previous empirical results also indicated the primary and extrastriatal visual cortex as the most consistently reported source, especially of high-frequency SSVEP (Duecker et al., 2020; Fawcett et al., 2004; Pan et al., 2021; Pastor et al., 2003; Vialatte et al., 2010; Zhigalov et al., 2019).

### Power versus coherence

When assessing the brain response to RFT stimulation, we also compared coherence with oscillatory power. Importantly, induced power provides estimates for each trial if required and so unlike coherence can provide the means for trial-based analysis, e.g. phase-to-amplitude coupling. Our results demonstrate that whilst oscillatory power does show a significant RFT effect across many conditions, it is a considerably less sensitive measure than coherence. This is explained by the fact, that the coherence estimate is based on the brain response phase-locked to the flicker thus reducing other noise contributions. In addition, there was one participant whose RFT response was extremely high as compared to others when measured with oscillatory power, especially in the RFT-FREQ session. The neurophysiological reasons for such high power values are unclear, but the analysis process needs to address such outliers. Coherence is less prone to such considerable differences amongst participants. Of course, extreme outlier values are not necessarily just a nuisance during the analysis, but could also reveal important neurophysiological and neurological health phenomena. When outlier values are not the focus of the research, we would recommend applying coherence or in the absence of photodiode evoked-power estimates (assuming the flicker input is phase-locked over trials; results not discussed) i.e. power estimates calculated after averaging across trials.

### Planar magnetometers versus magnetometers

The MEGIN MEG system contains planar gradiometer and magnetometer sensors with different signal-to-noise ratios and depth-sensitivities. The here analysed dataset provides the opportunity to assess and compare the results of the two sensor-types. The sensor-level findings were highly comparable for the two sensor-types and the source localization also indicated similar sources of the RFT effects; in most conditions around 10mm differences between the source maxima of the magnetometers and planar gradiometers. Yet the two sensors showed some differences. Firstly, the magnetometers compared to planar gradiometers showed higher mean coherences over selected posterior sensors and also showed significant RFT effects even for smaller stimulation patch-sizes and at more positions. This suggests a generally higher sensitivity of the magnetometers to RFT. Secondly, whilst there was considerable agreement on the sensor-level and source-level results, the findings also indicate an important difference in terms of lateralization due to RFT area position being in the left or the right hemifield. Whereas the planar gradiometer results showed only one positive cluster in one condition, the magnetometers revealed that at least in one comparison there was a clear double-dissociation between left and right hemifield presentation where the planar gradiometers show only weak or no sign of lateralization.

Lastly, it has previously been reported that photic stimulation results in effects not only at the stimulation frequencies but also at harmonic frequencies due to a non-sinusoidal SSVEP (Pastor et al., 2007; Rager & Singer, 1998) and also subharmonics for reasons less clear (Herrmann, 2001). The planar gradiometer results with coherence in our study revealed RFT effects only at the stimulation frequency and no harmonics and sub-harmonic effects. In contrast, there was a small effect at the 2^nd^ harmonic frequency with 60Hz and 64Hz stimulation when the magnetometer data were analyzed in the same way, again showing an important difference between the conclusions drawn based on the two sensor-types.

These findings indicate that the magnetometer sensor type seems to be more sensitive to the RFT response. In a relatively noise-free environment it seems SNR is superior to gradiometer sensors and thus it is recommended to take the magnetometer signal into account when investigating cortical responses to such visual stimulation.

## Conclusion

With this extensive empirical investigation of invisible fast rhythmic stimulation, i.e. rapid frequency tagging (RFT), we propose a set of basic parameters which, if adhered to, will give optimal results when using RFT in cognitive neuroscience investigations. Specifically, we suggest a maximum tagging frequency of 72Hz and visual angle of approximately 9° or above as well as stimulation in the lower visual field. Moreover, analysing the coherence between the recorded brain signal with the actual stimulation signal proves to be more sensitive than power estimates. Finally, in all three experimental sessions (frequency, patch size and location) magnetometer sensors were more sensitive to the flicker signal compared to planar gradiometers.

## Author Contributions

TM and OJ conceived the study; TM and BB implemented and recorded the experiment; TM and BB analysed the data; TM, BB and OJ wrote the manuscript

