## Supplementary Figures for "Optimal parameters for Rapid Invisible Frequency Tagging using MEG"

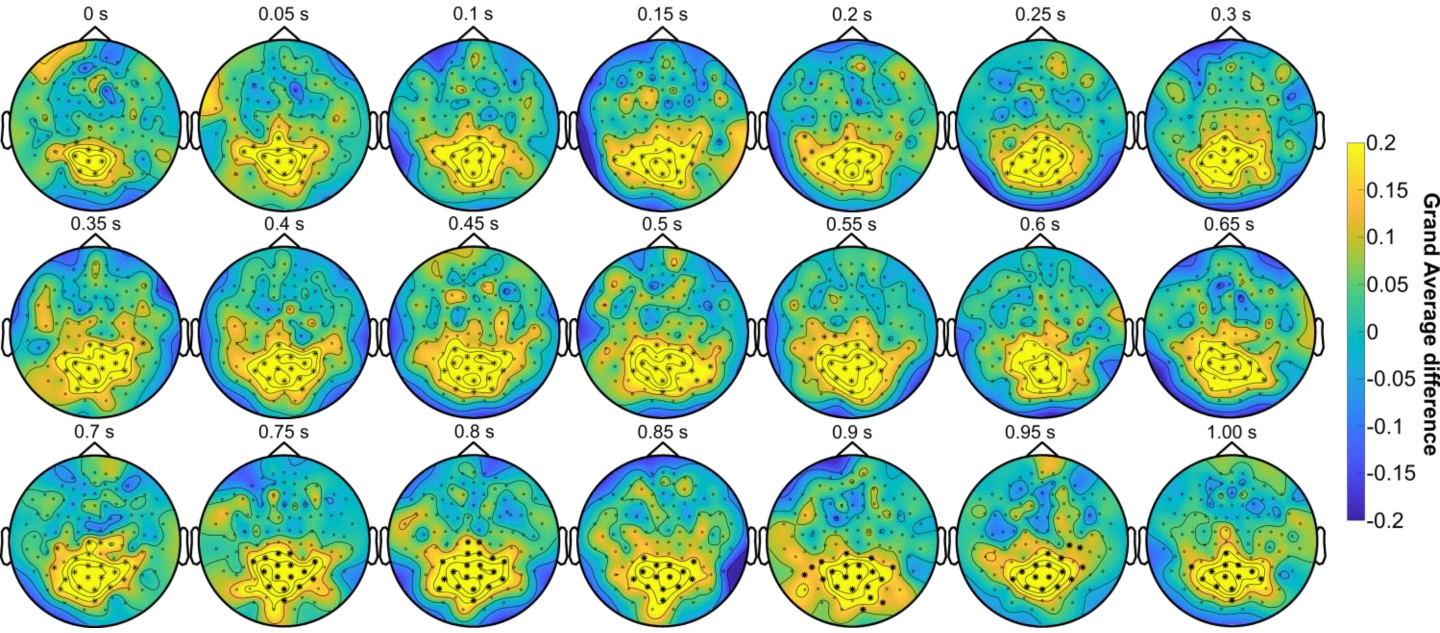


Figure S1. Topography of the grand average ***coherence*** difference between RFT and baseline time-windows at 60Hz in response to 60Hz stimulation over time. Significant differences ($p<.05$) between the RFT and baseline time-windows are shown over highlighted sensors (cluster permutation test). **Planar gradiometer** results.

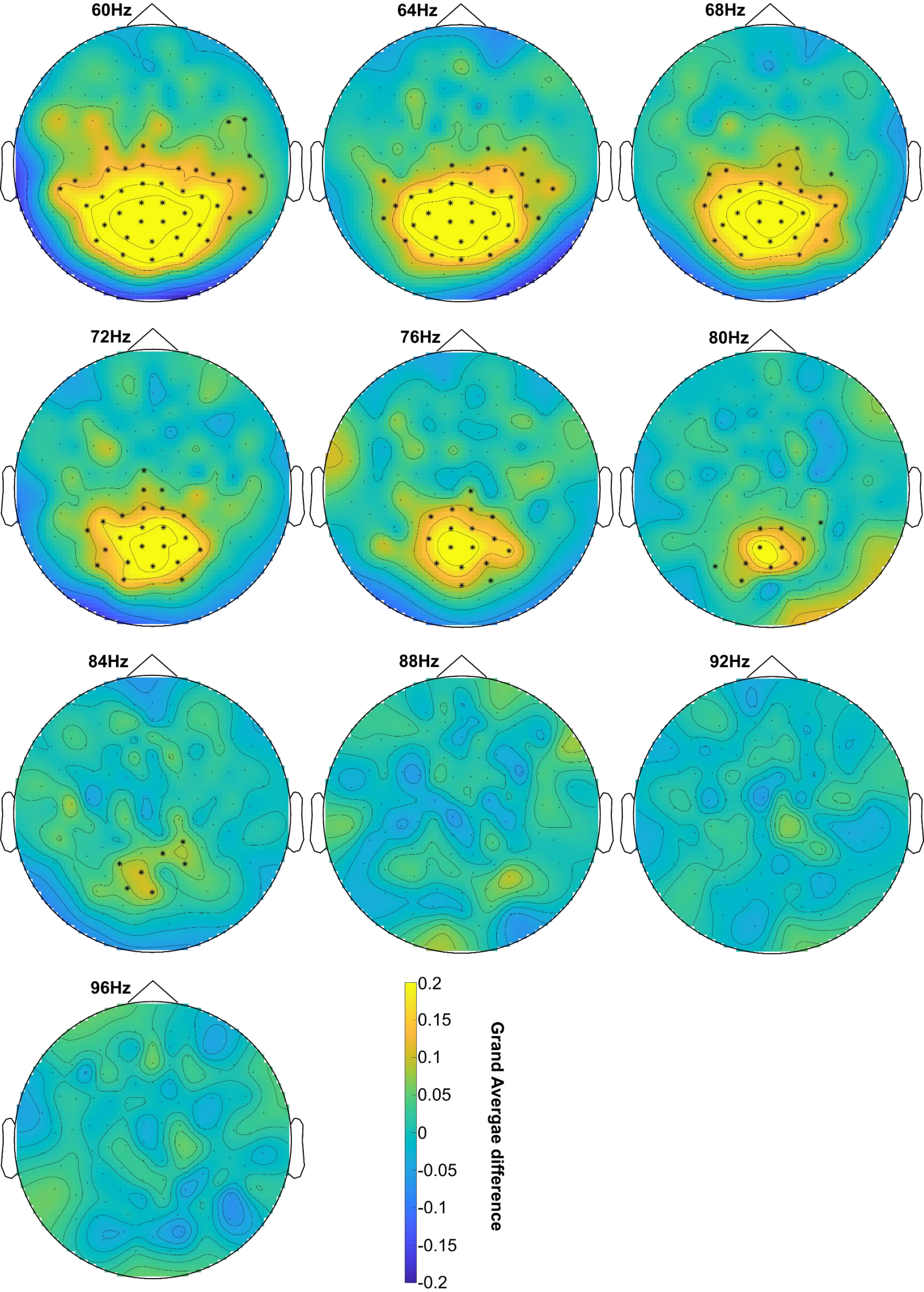


Figure S2. Topography of the grand average ***coherence*** difference between RFT and baseline time-windows at frequencies of 60Hz to 96Hz in 4Hz steps in response to RFT at the corresponding stimulation frequency. Significant differences (p<.05) between the RFT and baseline time-windows are shown over highlighted sensors (cluster permutation test). **Planar gradiometer** results.

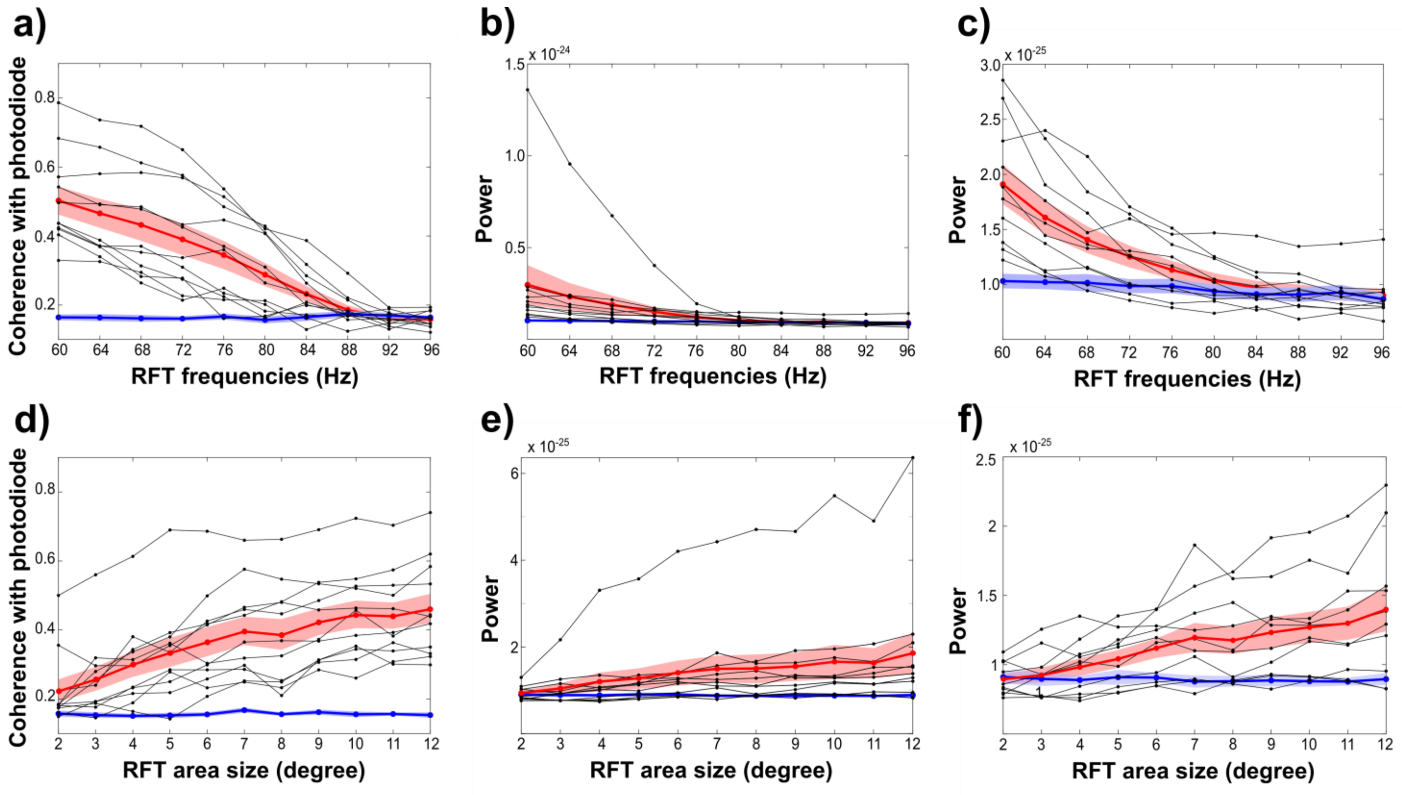


Figure S3. Grand average RFT (red line) and baseline (blue line) ***coherence*** as well as ***power*** over selected posterior sensor cluster. Shaded area indicates Standard Error of Mean. Grey lines show individual coherence and power. (a) RFT-FREQ coherence. (b) RFT-FREQ power with one outlier. (c) RFT-FREQ power without outlier. (d) RFT-SIZE coherence. (e) RFT-SIZE power with one outlier. (f) RFT-SIZE power without outlier. **Planar gradiometer** results.

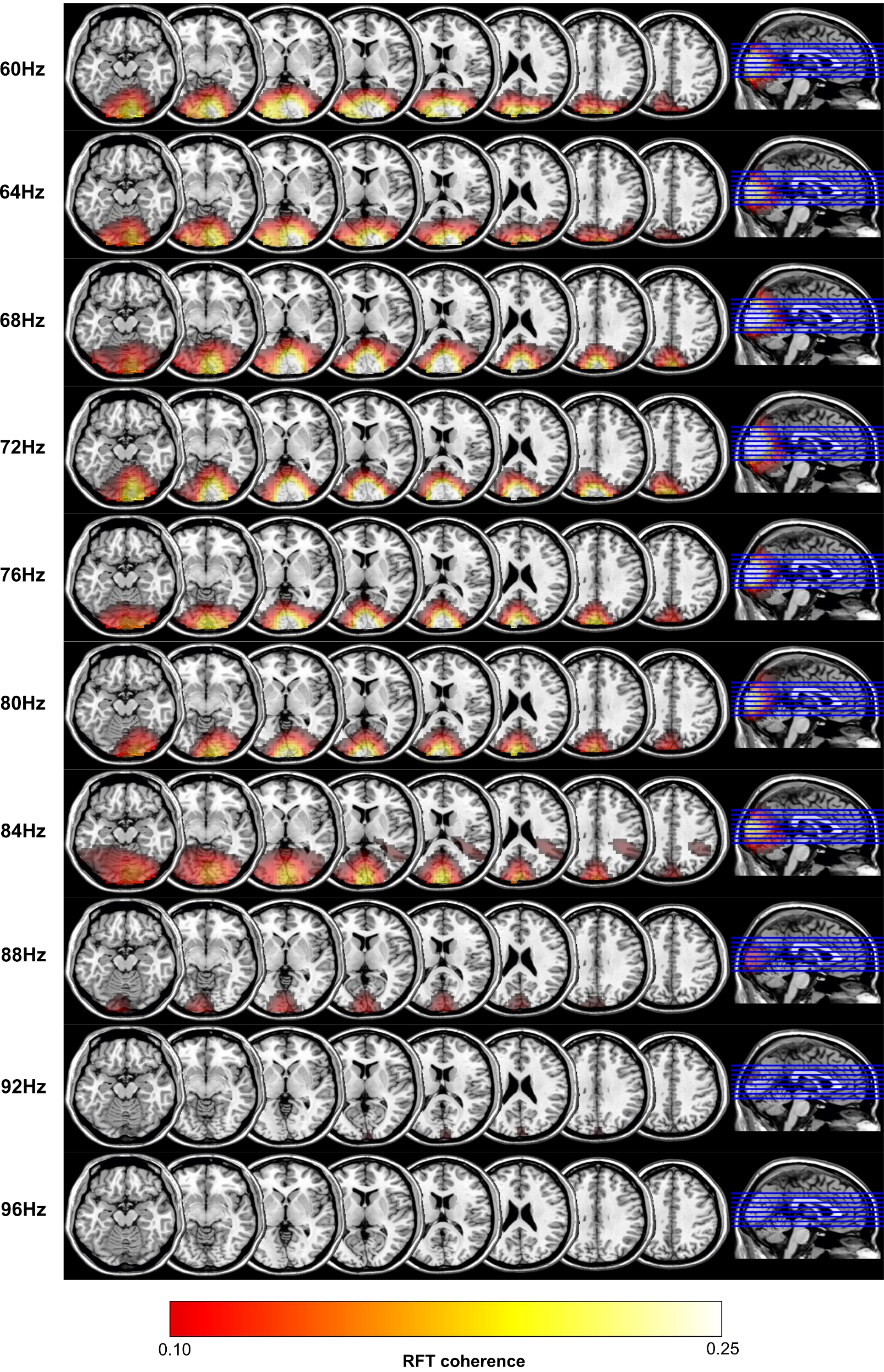


Figure S4. Grand average of the source localization results with DICS beamforming of the ***coherence*** in the RFT stimulation time-window at frequencies of 60Hz to 96Hz in 4Hz steps in response to the corresponding frequency stimulation. Results projected onto a MNI template brain. **Planar gradiometer** results.

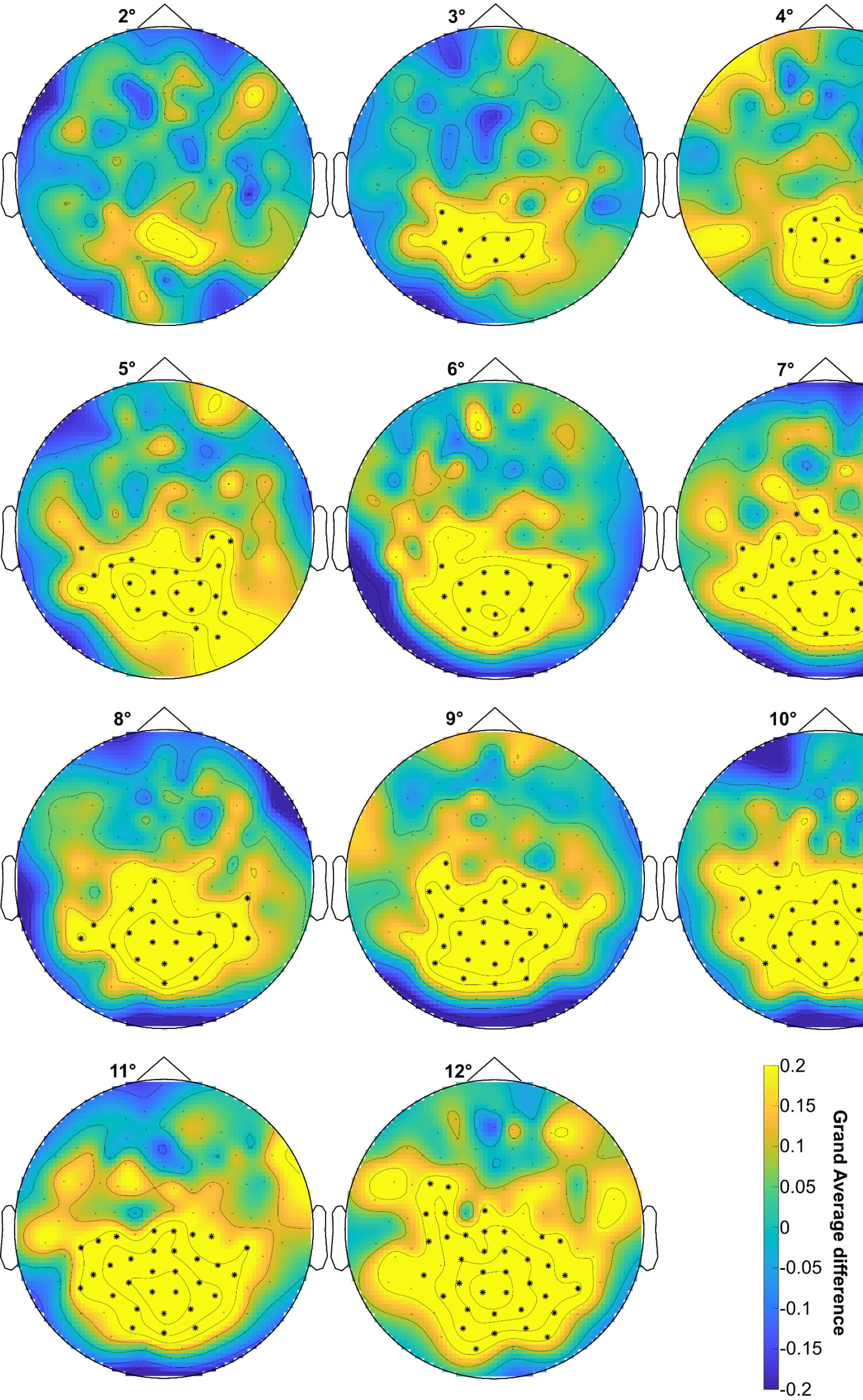


Figure S5. Topography of the grand average ***coherence*** difference between RFT and baseline time-windows at 66Hz in response to RFT at 66Hz with RFT area size from 2° to 12°. Significant differences ($p<.05$) between the RFT and baseline time-windows are shown over highlighted sensors (cluster permutation test). **Planar gradiometer** results.

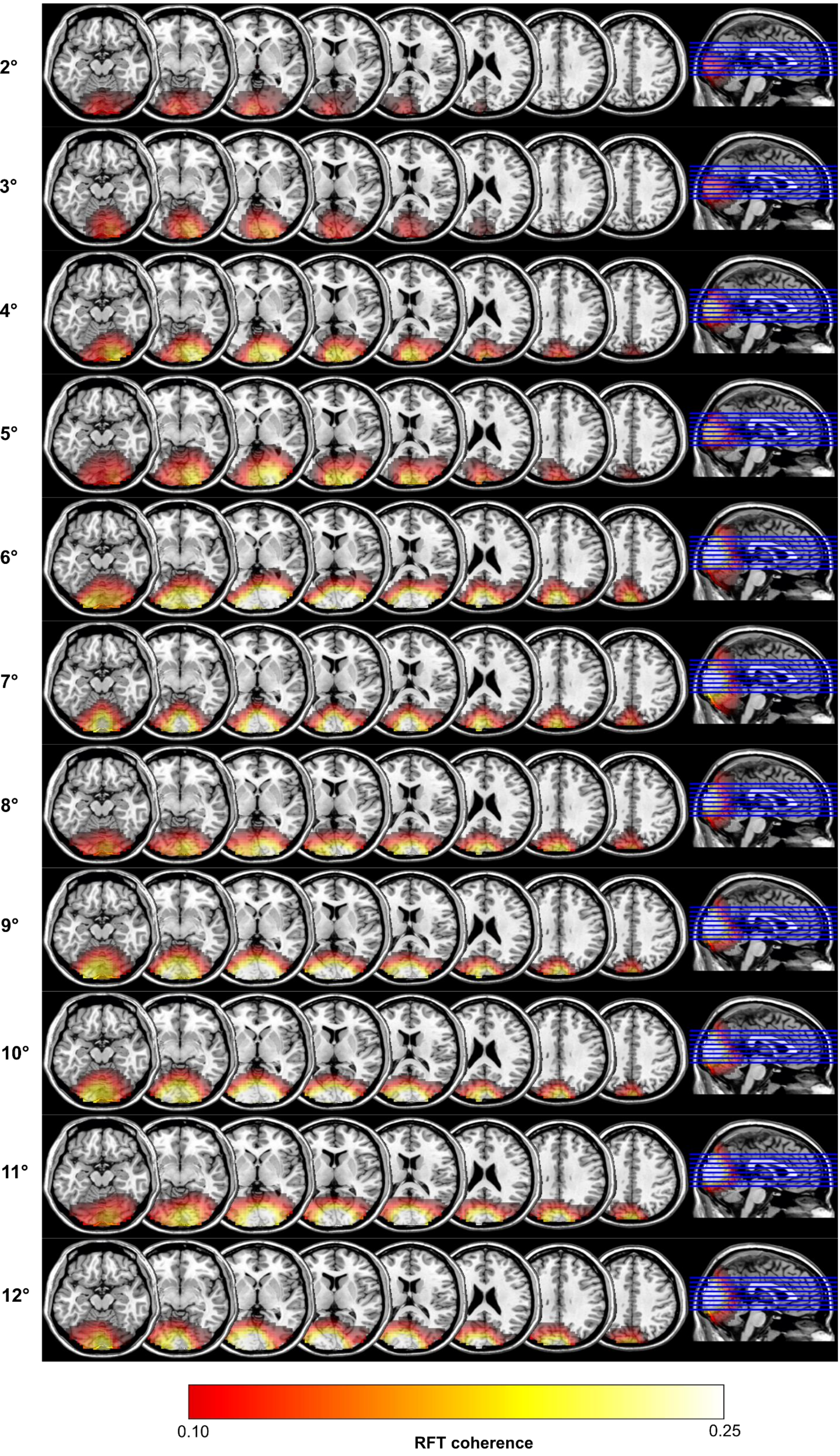


Figure S6. Grand average of the source localization results with DICS beamforming of the ***coherence*** in the RFT stimulation time-window at 66Hz in response to 66Hz stimulation with RFT area size from 2° to 12°. Results projected onto a MNI template brain. **Planar gradiometer** results.

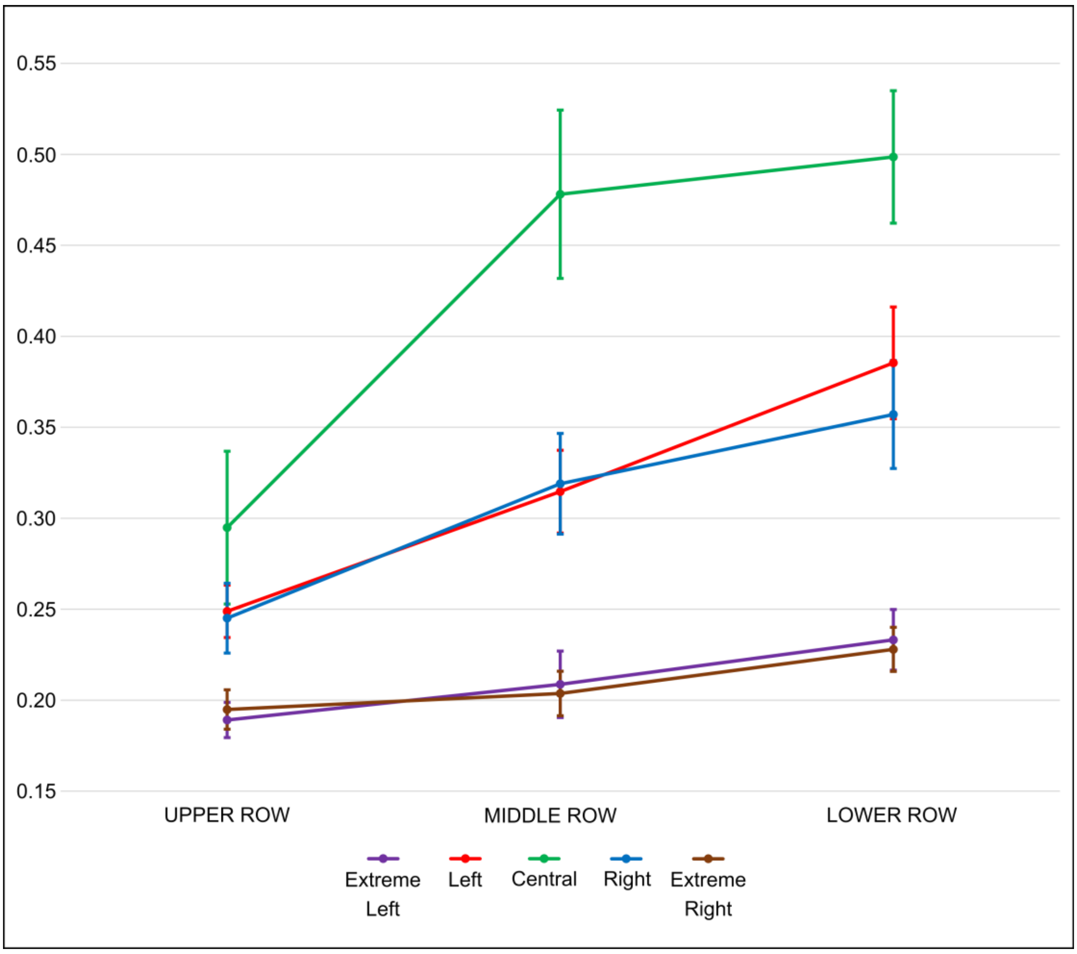


Figure S7. Grand average ***coherence*** over selected posterior sensors in response to RFT delivered at different positions on the screen. **Planar gradiometer** results.

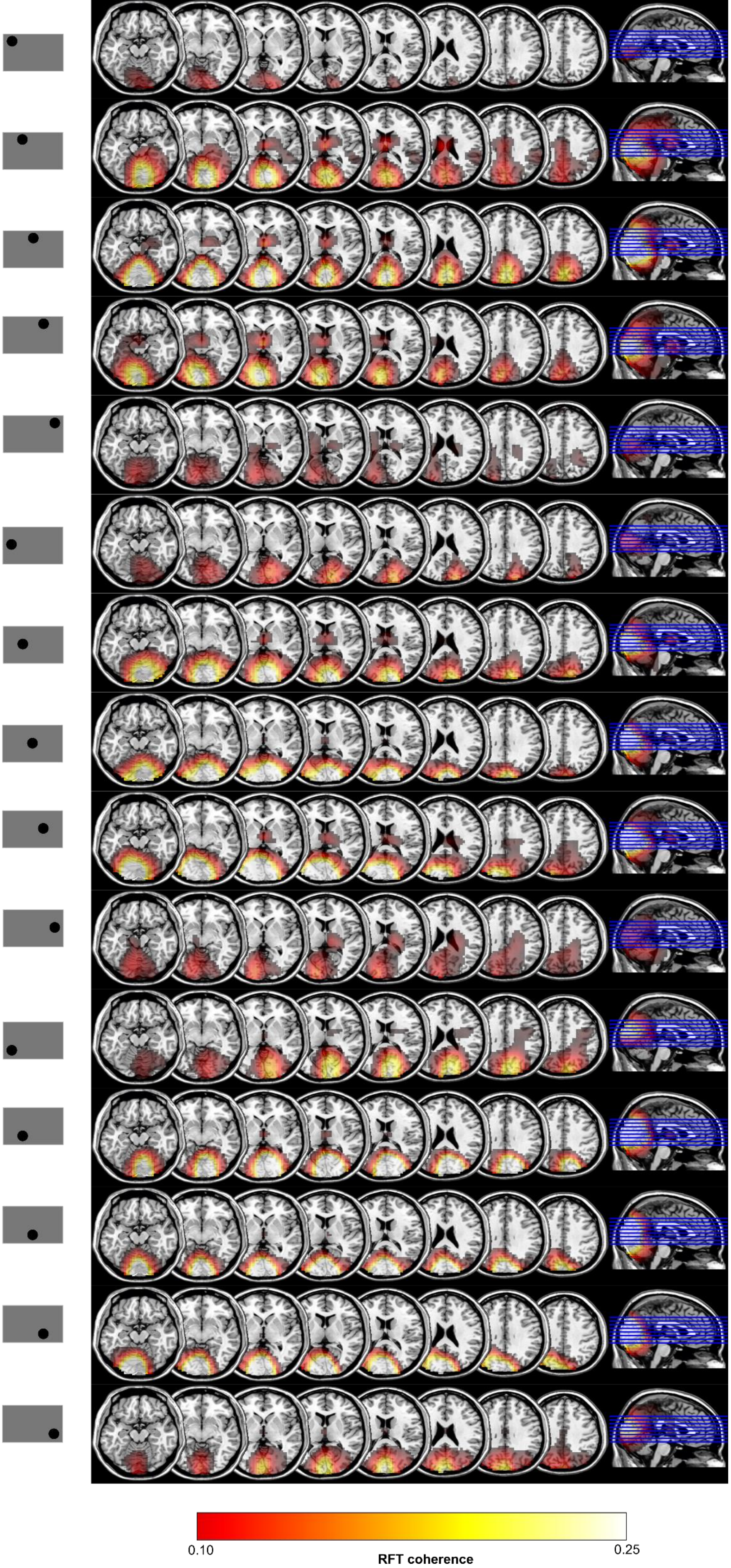


Figure S8. Grand average of the source localization results with DICS beamforming of the ***coherence*** in the RFT stimulation time-window at 66Hz in response to 66Hz stimulation and different RFT positions at the screen. Results projected onto a MNI template brain. **Planar gradiometer** results.

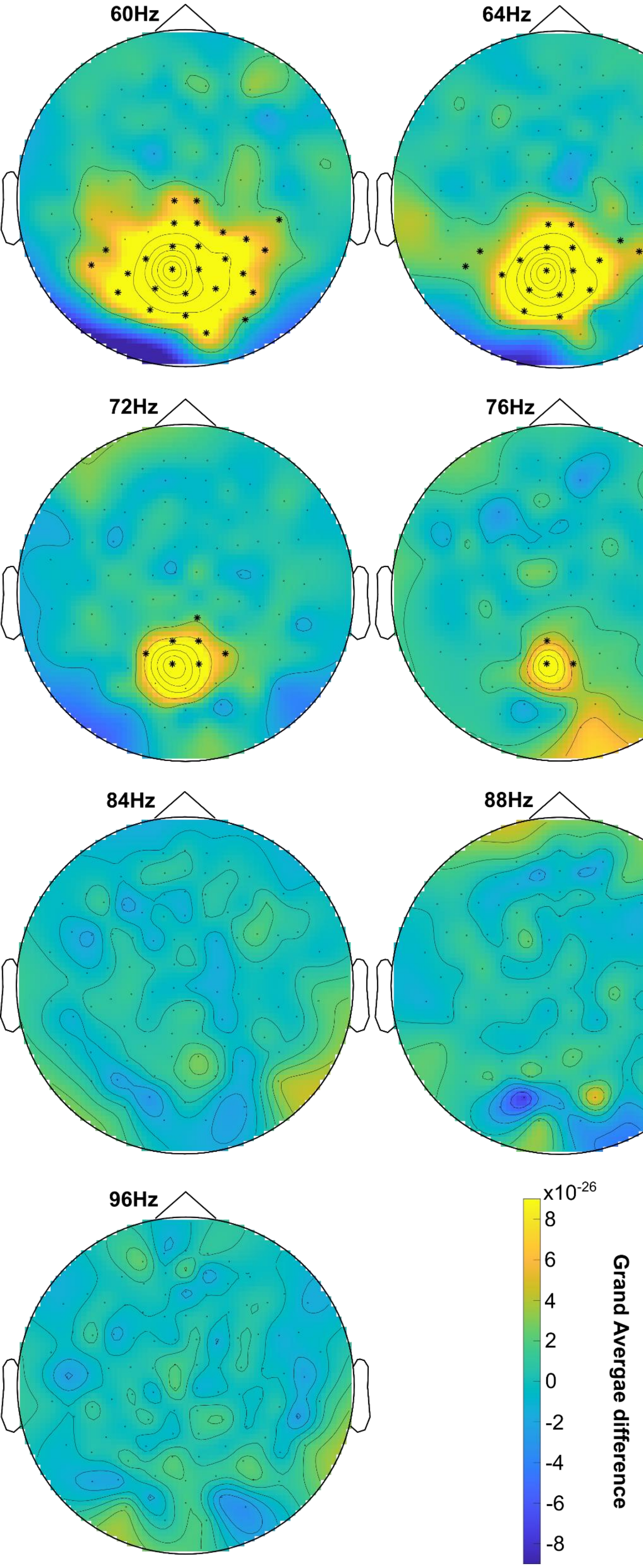


Figure S9. Topography of the grand average ***power*** difference between RFT and baseline time-windows at 60Hz to 96Hz in 4Hz steps in response to RFT at the corresponding stimulation frequency. Significant difference ($p<.05$) between the RFT and baseline time-windows are shown over highlighted sensors (cluster permutation test). **Planar gradiometer** results.

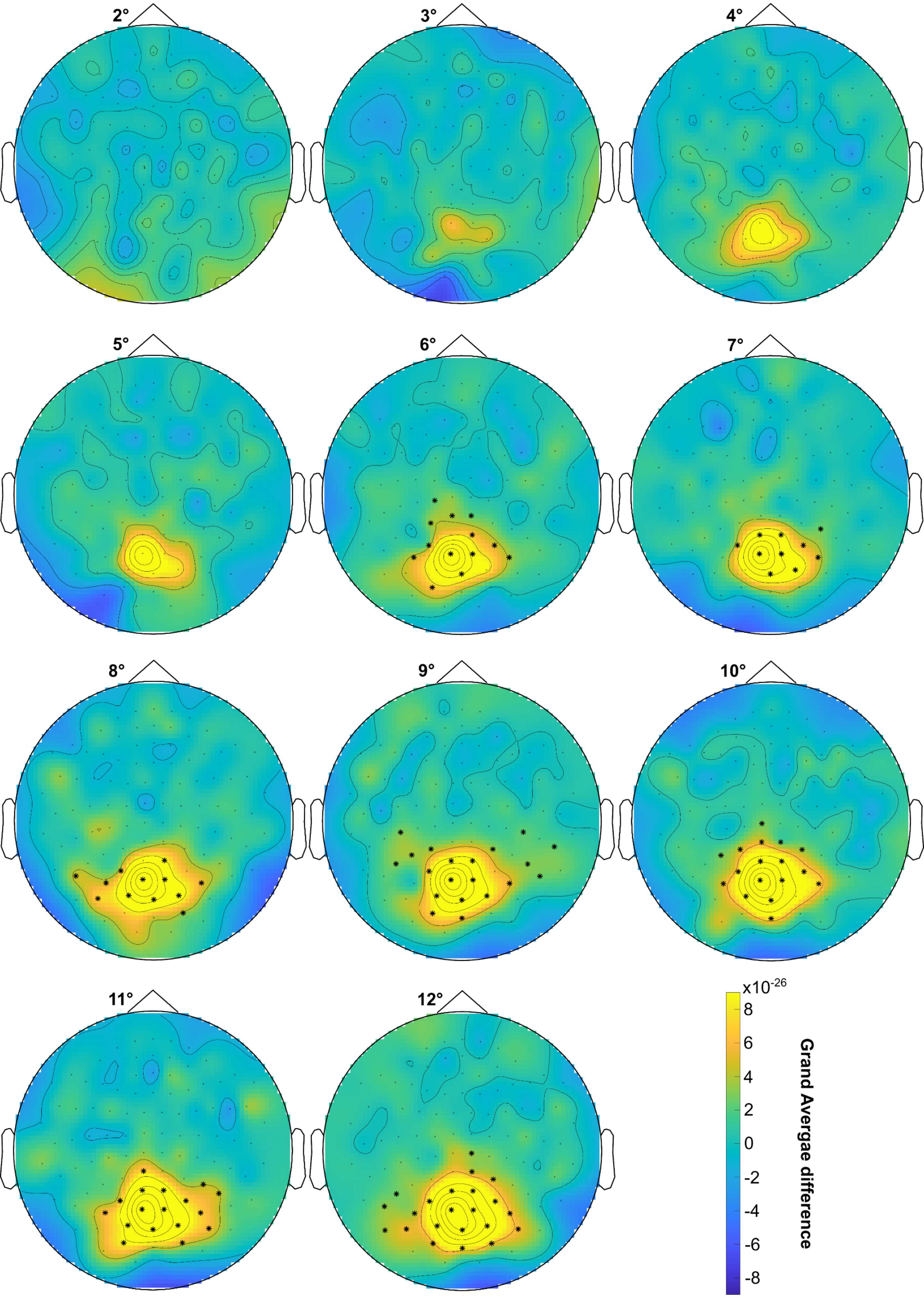


Figure S10. Topography of the grand average ***power*** difference between RFT and baseline time-windows at 66Hz in response to RFT at 66Hz with RFT area size from 2° to 12°. Significant differences (p<.05) between the RFT and baseline time-windows are shown over highlighted sensors (cluster permutation test). **Planar gradiometer** results.

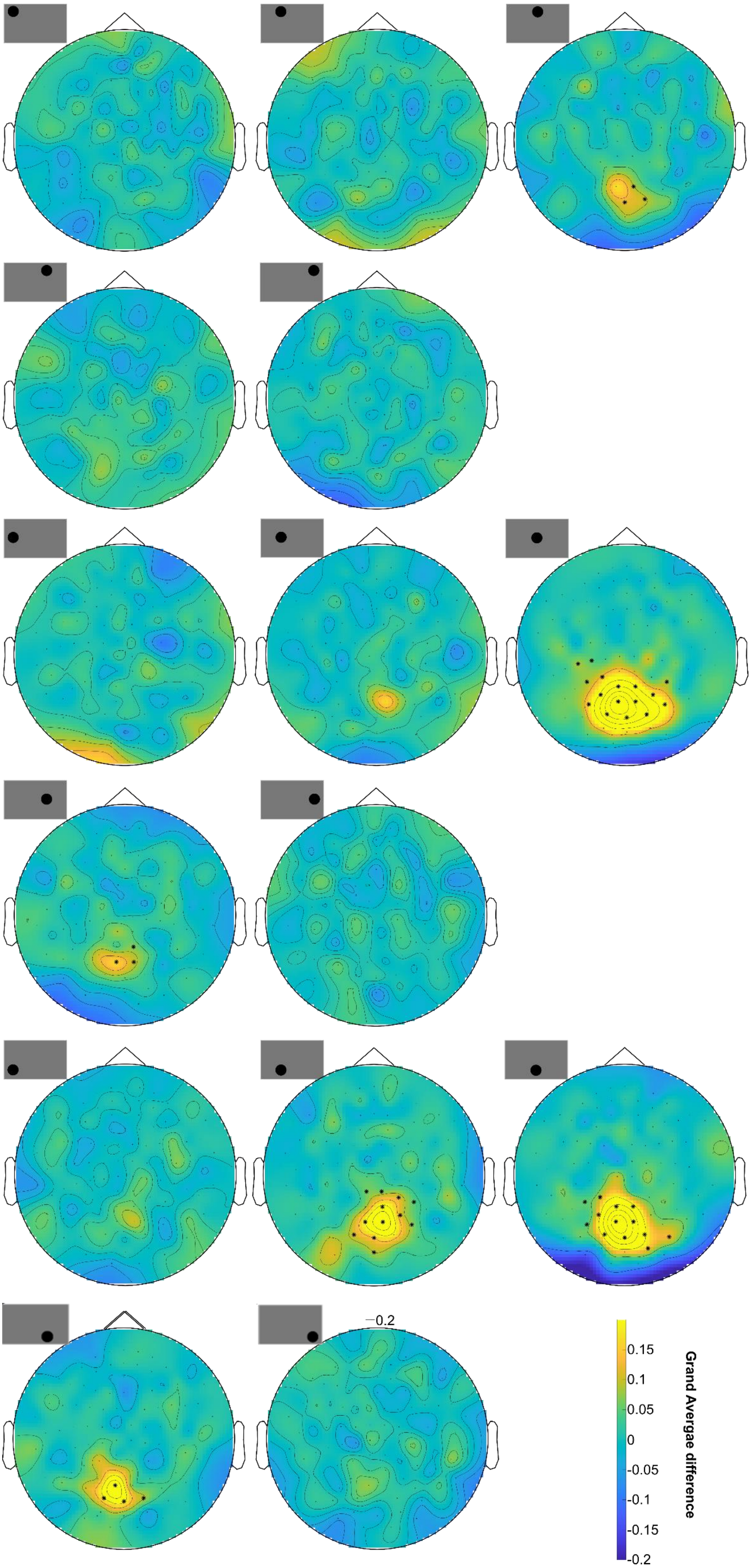


Figure S11. Topography of the grand average ***power*** difference between RFT and baseline time-windows at 66Hz in response to RFT at 66Hz and different positions at the screen. Significant differences (p<.05) between the RFT and baseline time-windows are shown over highlighted sensors (cluster permutation test). **Planar gradiometer** results.

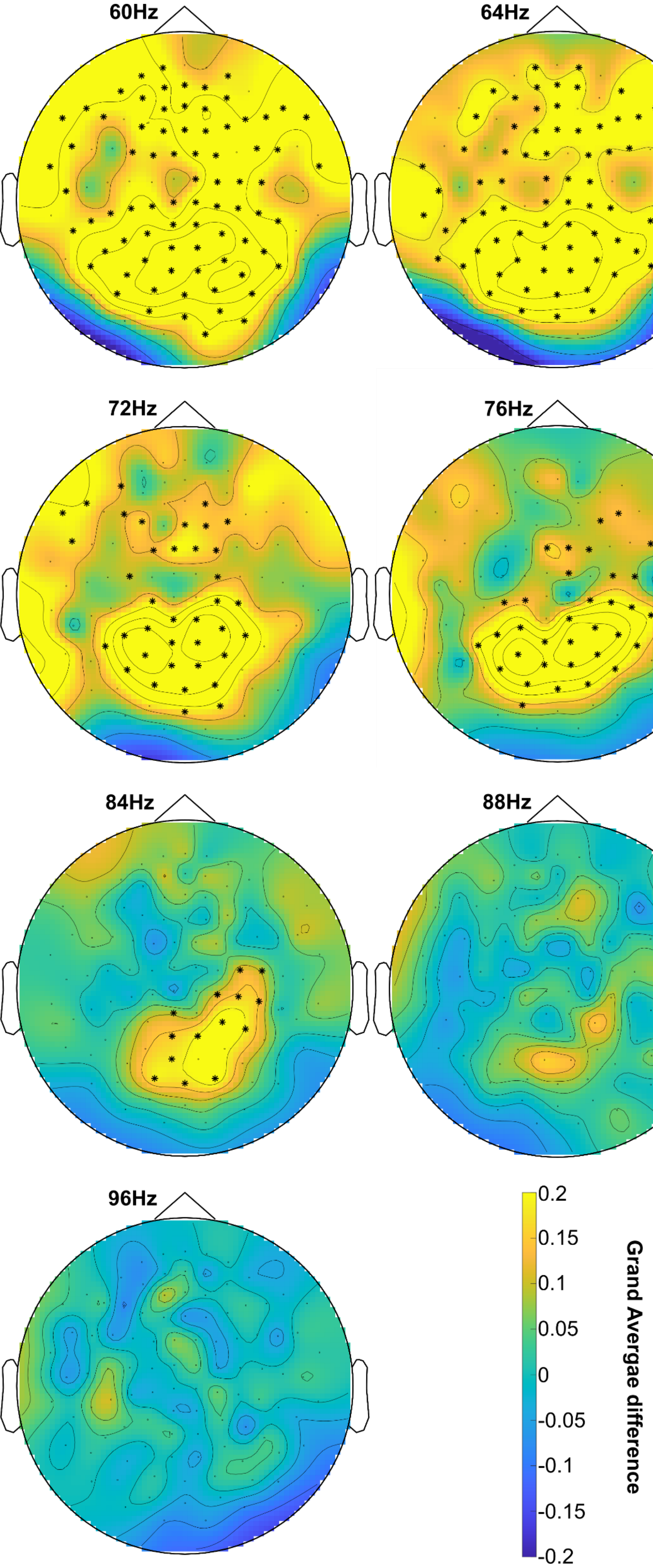


Figure S12. Topography of the grand average ***coherence*** difference between RFT and baseline time-windows at frequencies of 60Hz to 96Hz in 4Hz steps in response to RFT at the corresponding stimulation frequency. Significant differences (p<.05) between the RFT and baseline time-windows are shown over highlighted sensors (cluster permutation test). **Magnetometer** results.

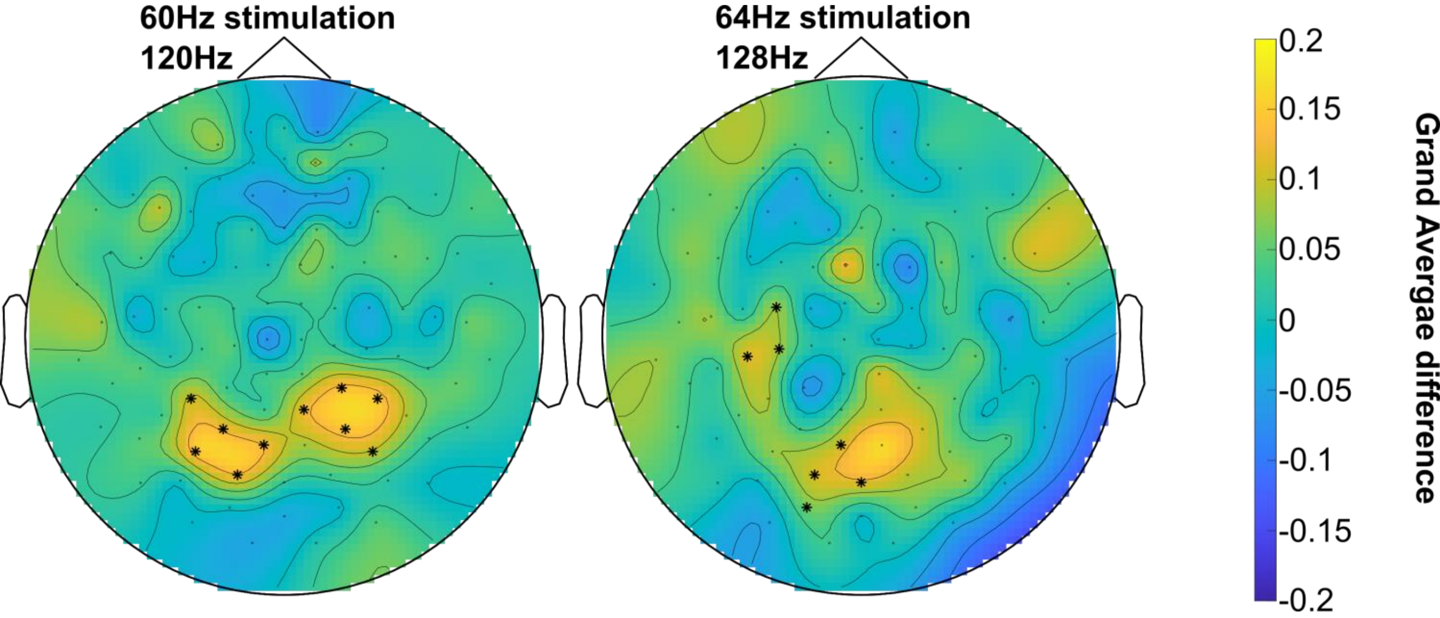


Figure S13. Topography of the grand average ***coherence*** difference between RFT and baseline time-windows at 120Hz and 128 Hz in response to RFT at 60Hz and 64Hz stimulation. Significant differences (p<.05) between the RFT and baseline time-windows are shown over highlighted sensors (cluster permutation test). **Magnetometer** results.

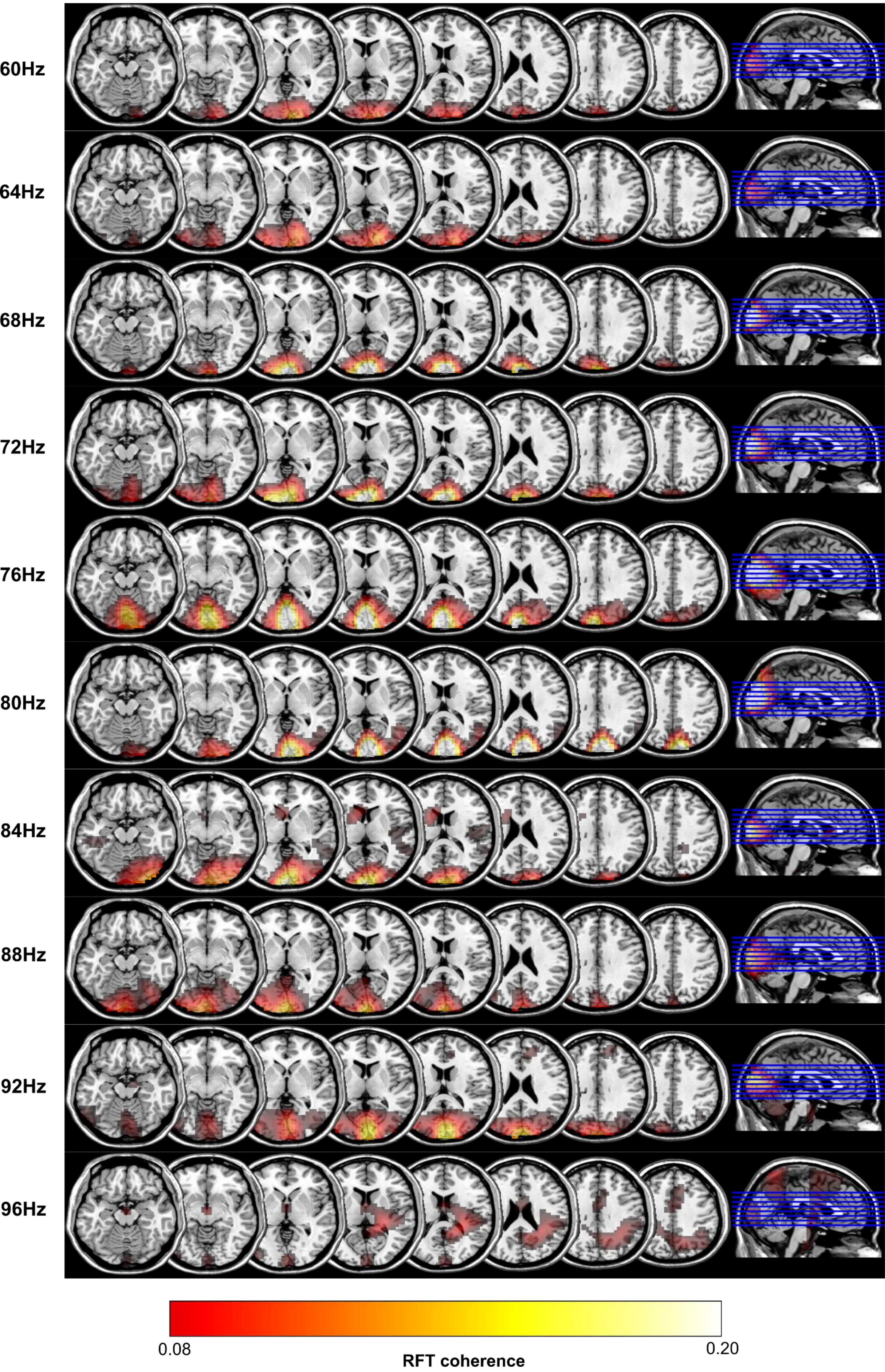


Figure S14. Grand average of the source localization results with DICS beamforming of the ***coherence*** in the RFT stimulation time-window at frequencies of 60Hz to 96Hz in 4Hz steps in response to the corresponding frequency stimulation. Results projected onto a MNI template brain. **Magnetometer** results.

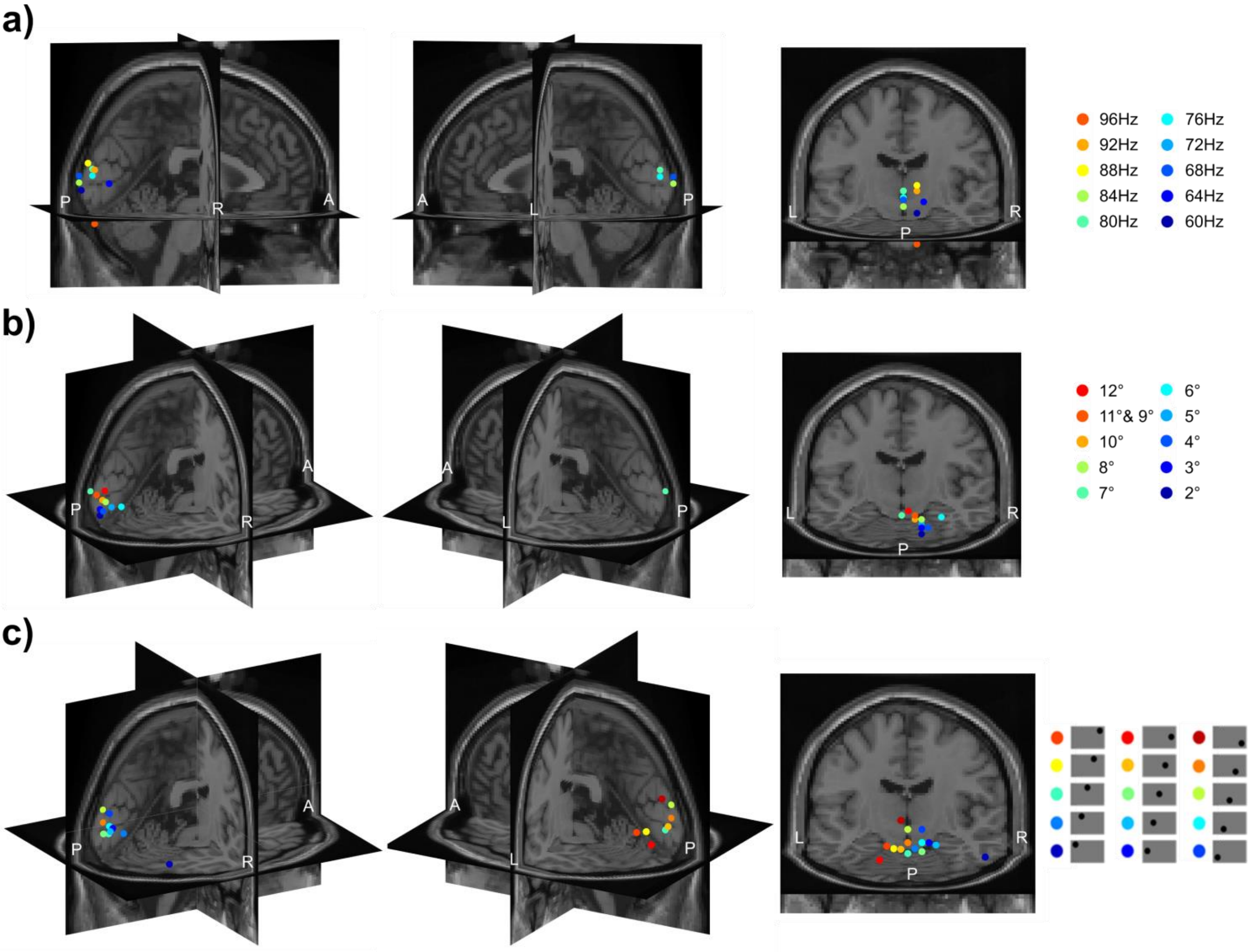


Figure S15. Maximum of the grand average of the source localization RFT ***coherence*** results with DICS beamforming of each condition in the RFT stimulation time-window projected onto a MNI template brain. (a) RFT-FREQ. (b) RFT-SIZE. (c) RFT-POSITION. **Magnetometer** results.

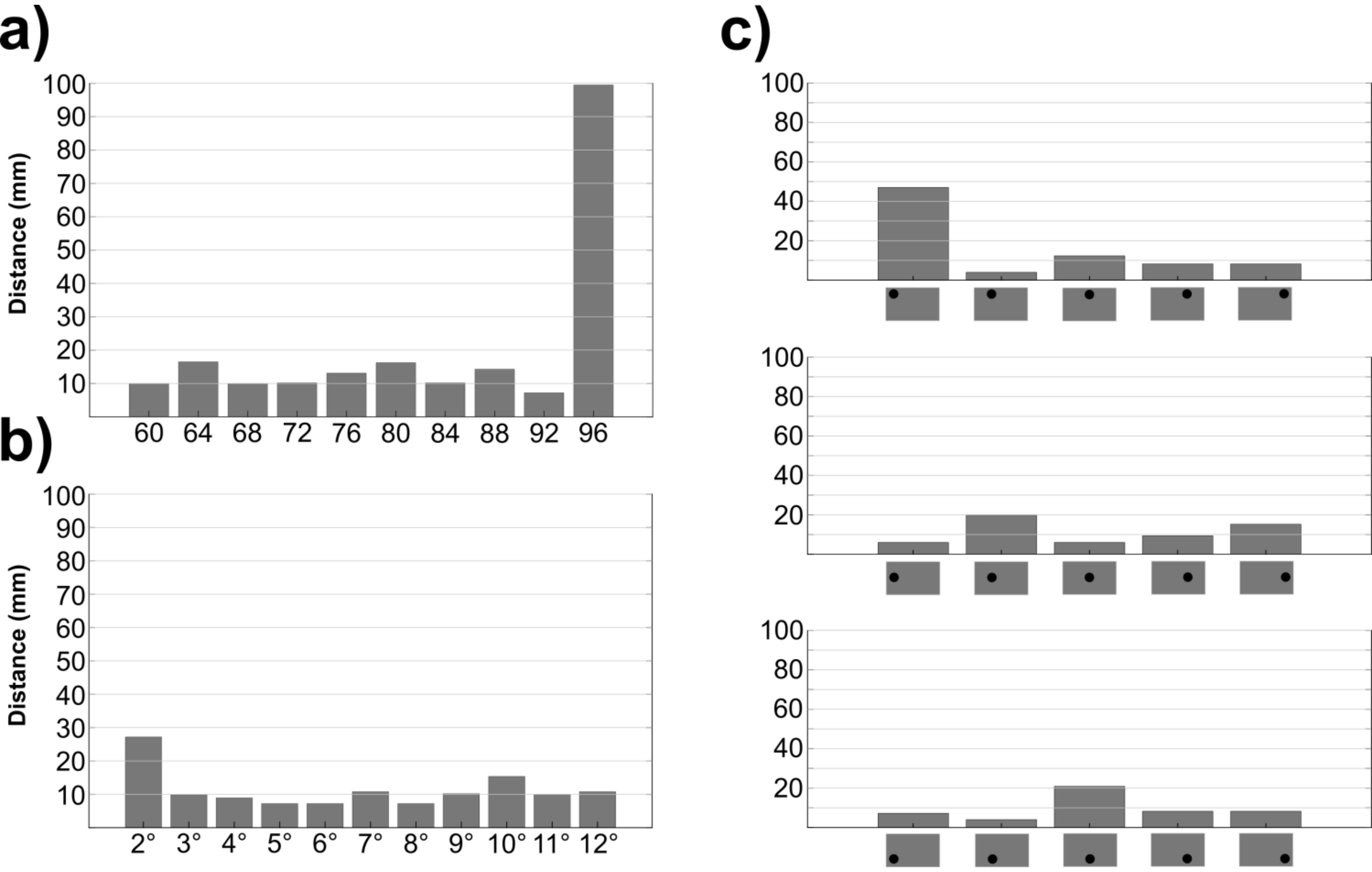


Figure S16. Maximum distance between the source maxima coordinates of the **planar gradiometer** and **magnetometer** ***coherence*** across the RFT conditions. (a) RFT-FREQ. (b) RFT-SIZE. (c) RFT-POSITION.

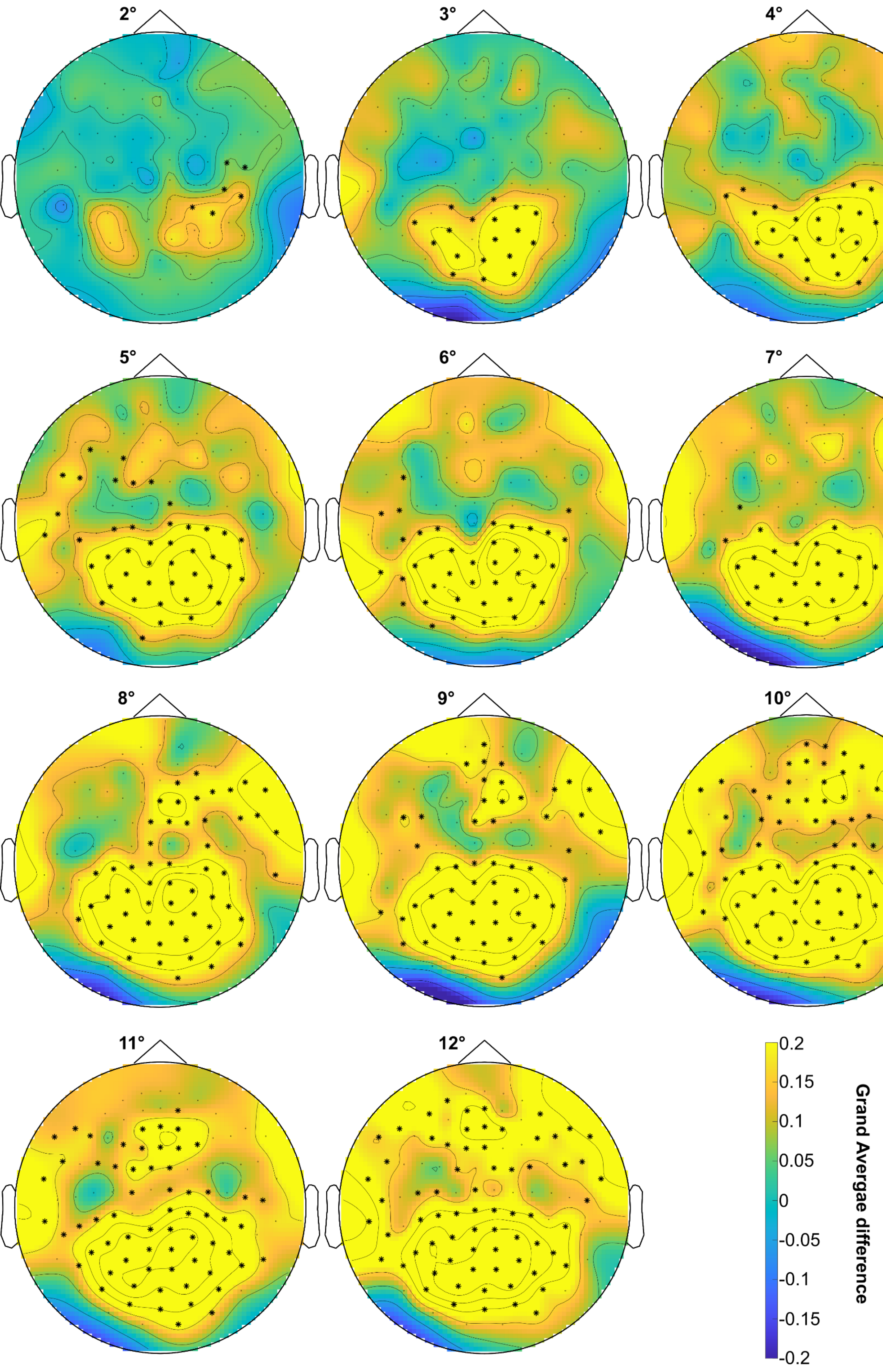


Figure S17. Topography of the grand average ***coherence*** difference between RFT and baseline time-windows at 66Hz in response to RFT at 66Hz with RFT area size from 2° to 12°. Significant differences ($p<.05$) between the RFT and baseline time-windows are shown over highlighted sensors (cluster permutation test). **Magnetometer** results.

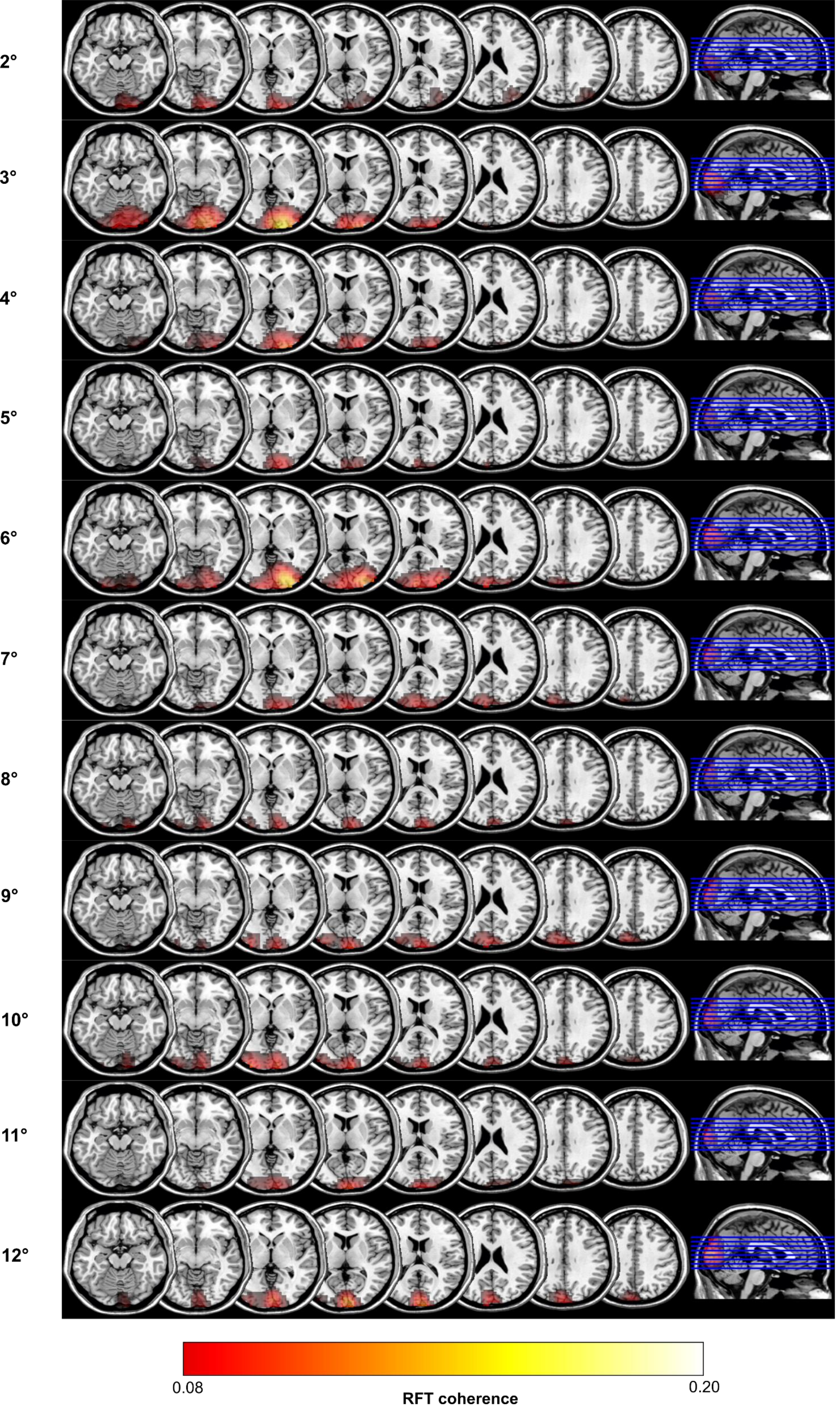


Figure S18. Grand average of the source localization results with DICS beamforming of the ***coherence*** in the RFT stimulation time-window at 66Hz in response to 66Hz stimulation with RFT area size from 2° to 12°. Results projected onto a MNI template brain.
**Magnetometer** results.

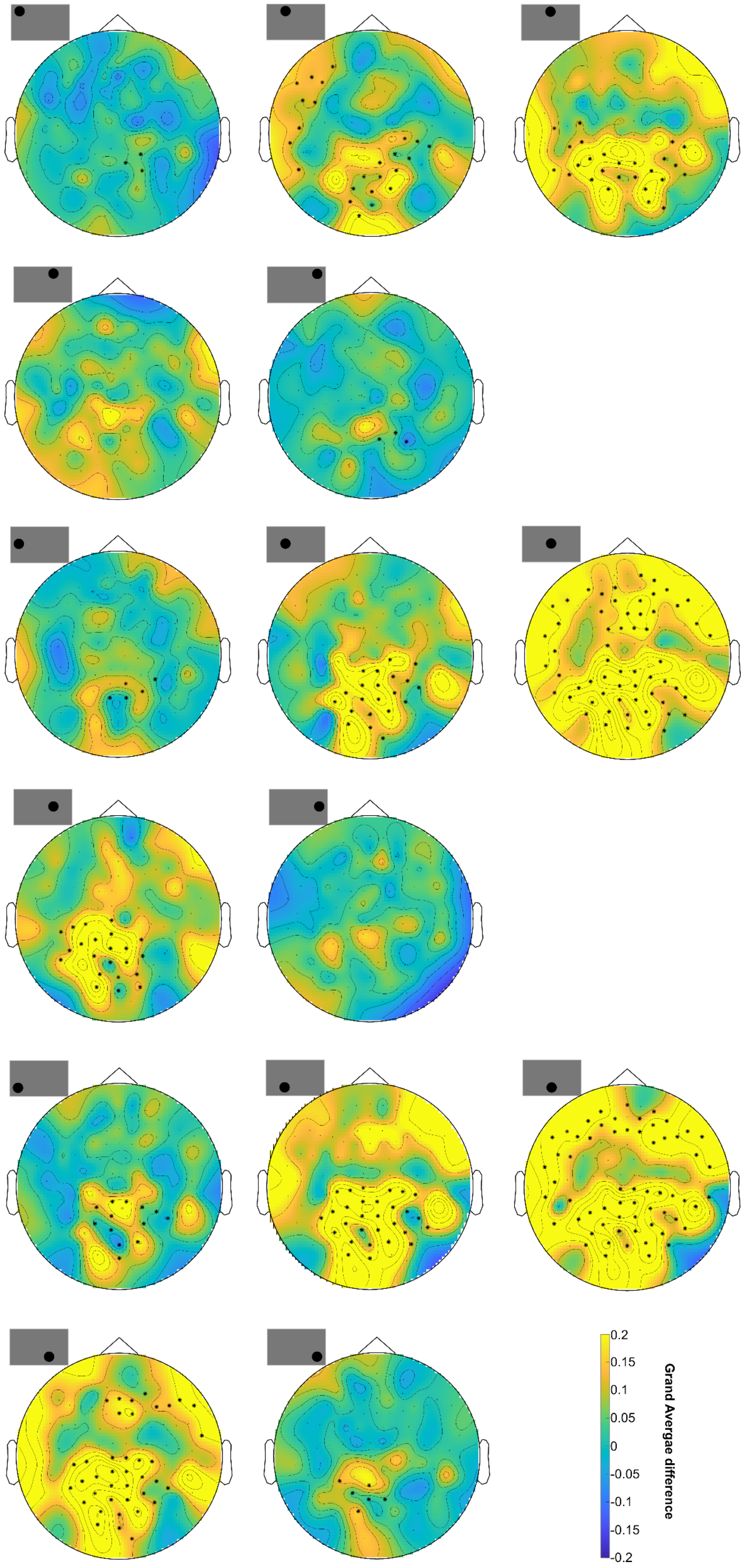


Figure S19. Topography of the grand average ***coherence*** difference between RFT and baseline time-windows at 66Hz in response to RFT at 66Hz and different positions at the screen. Significant differences (p<.05) between the RFT and baseline time-windows are shown over highlighted sensors (cluster permutation test). **Magnetometer** results.

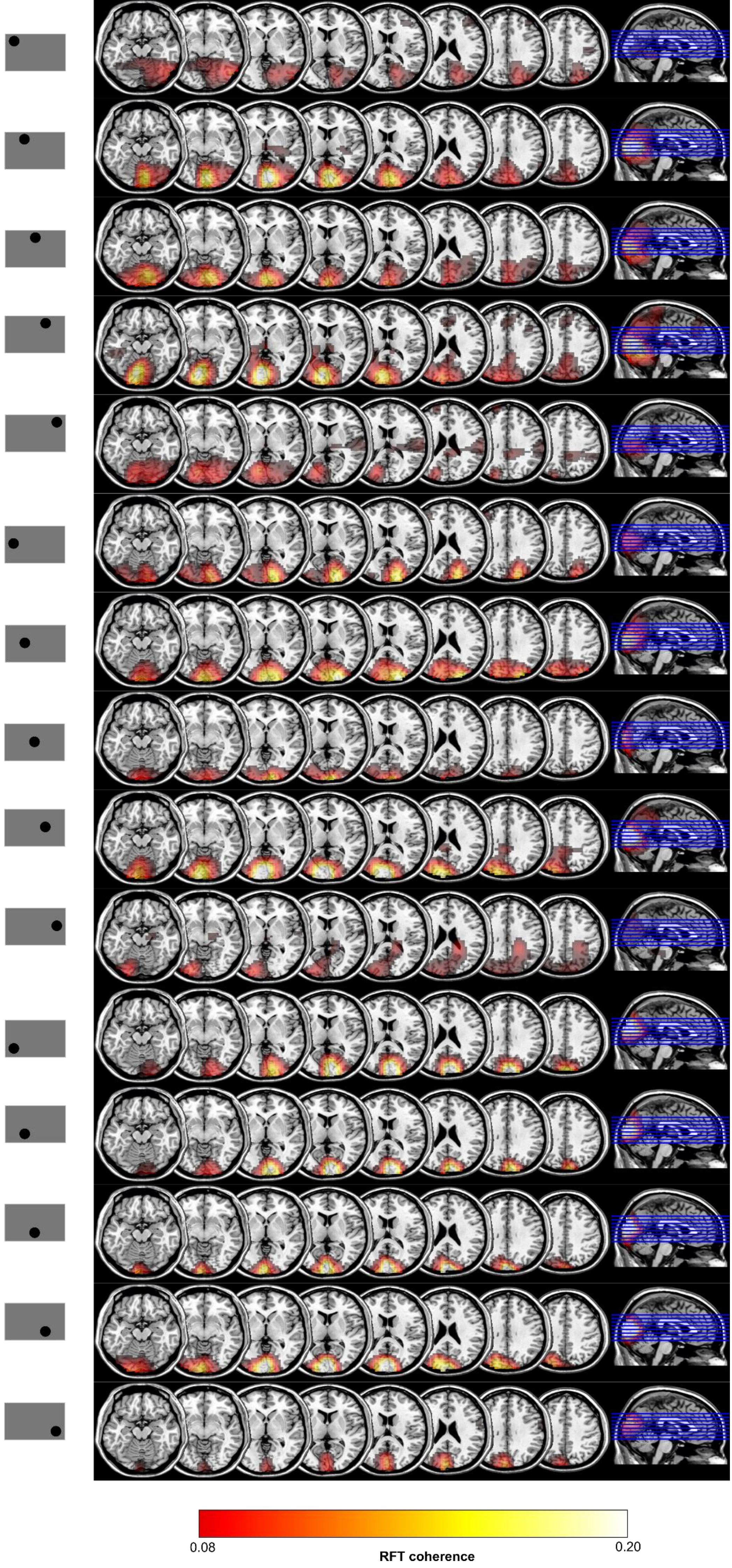


Figure S20. Grand average of the source localization results with DICS beamforming of the ***coherence*** in the RFT stimulation time-window at 66Hz in response to 66Hz stimulation and different RFT positions at the screen. Results projected onto a MNI template brain. **Magnetometer** results.
